# The coordination between CTCF, cohesin and TFs impacts nucleosome repositioning and chromatin insulation to define state specific 3D chromatin folding

**DOI:** 10.1101/2024.11.02.620823

**Authors:** Catherine Do, Guimei Jiang, Giulia Cova, Paul Zappile, Adriana Heguy, Jane A Skok

**Affiliations:** Department of Pathology, NYU Grossman School of Medicine, New York, NY, USA; Perlmutter Cancer Center, NYU Langone Health, New York, NY, USA; Genome Technology Center, NYU Grossman School of Medicine, New York, NY, USA

## Abstract

CTCF-mediated chromatin folding plays a key role in gene regulation, however the mechanisms controlling chromatin organization across cell states are not fully elucidated. Comprehensive analyses reveal that CTCF binding stability and cohesin overlap in mice and humans, are regulated by species specific differences in CTCF binding site (CBS) accessibility and enrichment of motifs corresponding to expressed TFs. By analyzing TFs, we confirm the co-operativity and competitiveness of TF/CTCF binding, which we further validate by allele specific analysis of SNPs. TF motif enrichment at CTCF bound sites is determined by cell state-specific transcriptional programs, which either stabilize or destabilize CTCF binding, as reflected by changing TF concentration. To examine CTCF binding in the context of nucleosome positioning, we performed single molecule nano-NOMe-seq. Subsetting reveals a continuum of binding states for CTCF, which are differentially represented at accessible versus inaccessible CBSs. As expected, CTCF degradation leads to a progressive loss of binding and nucleosome repositioning giving profiles similar to CTCF free CBSs. We also observe a similar time dependent effect when the cohesin subcomponent, SSC1 is degraded although CTCF remains bound, indicating that cohesin mediates CTCF-associated nucleosome repositioning. Stratified analysis of CTCF signal strength and accessibility reveals that in the presence of cohesin, CTCF strength contributes to nucleosome repositioning and chromatin insulation independent of accessibility. However, cobound TFs can uncouple the relationship between signal strength and nucleosome repositioning, without affecting the connection between repositioning and insulation. These studies identify mechanisms underlying cell state-specific CTCF profiles, linked to local and long-range chromatin organization.

## Introduction

CTCF is an eleven zinc finger protein that makes contact with a 12-15bp DNA motif through its central ZFs, 3-7. Together CTCF and the cohesin complex promote the formation of chromatin loops within highly self-interacting topologically associated domain (TAD) structures, via a loop extrusion mechanism ^1,2^. The two factors are enriched at TAD boundaries where they function as insulators promoting interactions between enhancers and promoters within a domain, and restricting the action of regulatory elements to target genes outside a domain. CTCF mediated chromatin folding plays a key role in gene regulation as exemplified by numerous studies showing that disruption of insulated TAD boundaries by deletion of CTCF binding sites (CBSs) can alter gene expression leading to defects in developmental processes and cancer initiation as a result of illegitimate enhancer promoter contacts ^3–7^.

CTCF dependent chromatin interactions can be conserved, or cell-state specific ^8^, however the mechanisms underlying CTCF’s ability to act as a chromatin organizer in a developmental stage, or disease specific manner have not been fully elucidated. We and others have found that CTCF can bind both accessible and inaccessible sites, with differential functional outcomes ^9,10^. By analyzing the interplay between CTCF, cohesin and ATAC-seq data in CTCF WT and mutant expressing cells, we demonstrated that accessibility is a key determinant of CTCF binding stability, and CTCF signal strength in turn is the strongest predictor of cohesin overlap and chromatin insulation score ^9^. Specifically, when CTCF binds accessible sites, it has a stronger signal and is more efficient at blocking cohesin and participating in chromatin loop formation than CTCF at inaccessible sites ^9^. However, the relationship between accessibility and CTCF binding stability is not clear cut, because a subset of inaccessible CTCF bound sites have strong signals with cohesin overlap, and conversely, CTCF at accessible sites can exhibit weak signals devoid of cohesin. Furthermore, there are numerous accessible CBSs that remain unbound. Thus, other factors must positively and negatively contribute to binding.

While previous studies have identified a handful of transcription factor (TF) cofactors that stabilize CTCF binding in specific cell types, a comprehensive analysis exploring the role of TFs in cell type-specific binding is lacking ^11–16^. In this study, we demonstrate that DNA sequences found within a 35 bp window surrounding CBSs motifs corresponding to expressed cell type-specific TFs can differentially affect CTCF signal strength and its capacity to block cohesin, in a manner that depends on whether they are located in accessible or inaccessible chromatin. In both mice and humans, we identified enriched TF motifs showing strong similarity within specific windows, suggesting potential redundancy in TF-mediated regulation of CTCF binding and a mechanism for cell type-specific control. We also highlight differences between the two species: notably, mouse and human differ respectively in the proportion of inaccessible versus accessible CBSs in their genomes, which is reflected in the regulatory mechanisms that govern their binding. Although TF motifs are present at most binding sites in both species, in mice they are enriched at inaccessible sites, while in humans they are more abundant at accessible sites.

To examine the impact of transcriptional programming on CTCF binding, we looked at how variations in individual TF levels across different cell states and cell types are associated with changes in their motif enrichment at CTCF sites. In contrast to previous findings ^11–16^, we show that TFs can function to both stabilize and destabilize CTCF binding, as reflected by the direction of enrichment at increasing TF concentration. By analyzing a handful of TFs, we could confirm both their co-operativity and competitiveness with CTCF binding at CBSs. Consistent with our ChIP-seq analysis, AlphaFold3 predicts that at individual locations, TFs can respectively compete or cobind with CTCF depending on whether TF binding overlaps the CBS or occurs up / downstream of it. Allele specific validation further highlights that SNPs affecting either CTCF or TF binding have reciprocal effects on the binding of each factor. Moreover, changes in binding of either factor can impact cohesin binding and to a lesser extent, ATAC-seq signals. Consequently, changes in TF expression levels can convert a weak CTCF-bound site into a strong site that halts the progression of cohesin, or vice versa, depending on whether the adjacent TF binds cooperatively or competitively with CTCF and whether its expression increases or decreases. This duality helps explain why many accessible CBSs are not occupied by CTCF itself, but rather by TFs that prevent its localization Using long read, nano-NOME-seq single molecule footprinting, we analyzed the impact of CTCF binding on nucleosome repositioning in different genomic contexts. The single molecule approach allowed us to subset our data revealing a continuum of binding states for CTCF which are present at individual peaks, and differentially represented at accessible and inaccessible sites. This information has been missing from previous bulk analyses that average all the data. Using the clustered analysis, we demonstrate that CTCF has a reduced ability to reposition nucleosomes at inaccessible sites. Furthermore, we found that (i) the profiles of degraded CTCF and CTCF free CBSs were very similar, reflecting nucleosome occupied CBSs or binding of TFs that can still phase nucleosomes but cannot reposition them, and (ii) degradation of the SCC1 subcomponent of cohesin gave rise to a similar profile while CTCF remains bound, pointing to the interesting finding that cohesin and loop extrusion mediate nucleosome positioning and phasing by creating or stabilizing the barrier effects at CTCF anchors.

A stratified analysis of CTCF signal strength and accessibility revealed that in the presence of cohesin, CTCF strength contributes to nucleosome repositioning and insulation independent of accessibility. However, cobound TFs can uncouple the relationship between signal strength and nucleosome repositioning without affecting the connection between nucleosome repositioning and insulation. These findings indicate that nucleosome repositioning is a more reliable predictor of chromatin insulation than CTCF signal strength and that TF cobinding can modulate loop extrusion at CTCF anchors. Together, these studies demonstrate that cell state-specific transcriptional programs, shape CTCF profiles linked to local and long-range chromatin organization and gene regulation.

## Results

### TF motifs enriched within 35 bps of CTCF bound sites stabilize CTCF binding

A number of studies have reported that CTCF binding can be stabilized by the cobinding of cell-type specific TFs adjacent to CTCF ^11–16^, but there have been no comprehensive, unbiased analyses to determine how cell type-specific cofactors regulate CTCF binding and its loop forming capacity. In addition, previous studies did not take into account the chromatin accessibility of CTCF binding sites, which can play an important role in CTCF cell-specific binding and function as previously shown by us and others^9,10^.

To investigate, we used mESC data previously generated by our lab ^9^ to examine the enrichment of TFs at defined windows in the 35 bp region surrounding the CTCF binding site (**Figure 1A**). We focused on this region as we observed little enrichment of TF motifs beyond this point (**Figure S1A**), although a slight increase could be detected further upstream between 50-75bps, that could impact CTCF binding via a different mechanism. In **Figure 1A** the x-axis shows the gap/distance in bps between the end of the TF motif and the start of the CTCF motif for upstream regions, or the start of the TF motif and the end of the CTCF motif for downstream regions. CTCF has eleven zinc fingers of which ZFs 3-7 make up the core binding domain that contacts the 12-15bp consensus motif. It has previously been reported that at some sites an Upstream (U) CTCF motif, can engage ZFs 9 to 11 and a less prevalent Downstream (D) motif can impair the binding of ZFs 1 and 2 to alter binding stability ^11,17–19^. We therefore defined CTCF upstream and downstream regions as the U motif window (U win) with binding ending at the 6 to 8 bp upstream gap, the U spacer as binding ending at the 0 to 7 bp upstream gap, the D spacer as binding beginning at the 0 to 9 bp downstream gap, and the D motif window (D win) as binding beginning at the 10 to 11 downstream gap.

**Figure 1:**
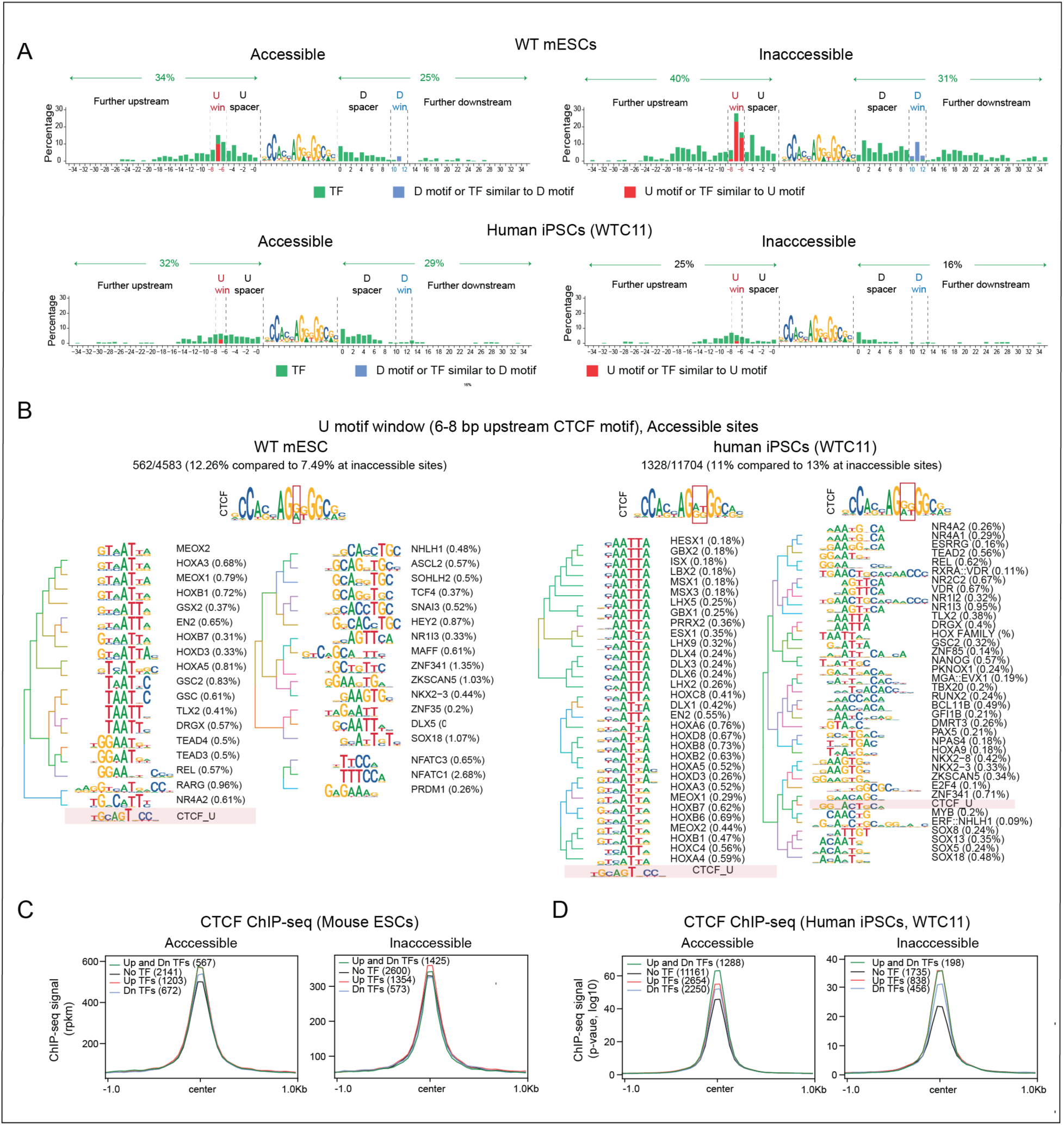
Bound CBSs are enriched in TF motifs within 35bps of the CTCF motif in mouse and human. A: Bar graphs showing the percentage of enriched TF motifs found using spacing motif enrichment analysis (SpaMo) at accessible and inaccessible sites in WT mESCs (top) and human iPSCs (bottom). The x-axis shows the gaps/distances in bp between the end of the TF motifs and the start of the CTCF motif for upstream gaps, or the start of the TF motif and the end of the CTCF motif for downstream gaps. For the U and D motif windows, the percentage of U and D motifs and TFs similar to the U and D motif are distinguished from other TF motifs (red or blue stacked bars, respectively). **B**: Motif logos of the enriched TF motifs found in the U motif window at accessible sites in mESCs (left) and human iPSCs (right). TF motifs ressembling the U motif and enrichment for the other windows are shown in **Figure S2-S6**,. **C-D**: CTCF binding profiles from ChIP-seq at accessible and inaccessible sites in mESCs (**C**) and human iPSCs (**D**) at CBSs containing at least one enriched TF motif within 35 bp upstream, downstream or both upstream and downstream compared to CBSs without any TF motifs.

### TF motifs have distinct enrichment profiles in mice and humans

Overall, in the mESCs, the percentage of TF motif enrichment (excluding U motifs and U like motifs) at both upstream and downstream regions highlights that TF motifs are predominantly associated with inaccessible (71%) rather than accessible (59%) sites (**Figure 1A**). A similar analysis in human WTC11 cells revealed that in contrast to mESCs, TF motifs are more enriched overall at accessible (61%) rather than inaccessible (41%) sites (**Figure 1A**). The differences between human and mouse could reflect the fact that there are proportionately more accessible CTCF binding sites in human compared to mouse (**Figure S1B** and **Table S1 A-B**).

### There is redundancy in the TFs that can bind a particular window

In **Figure 1A**, TFs with distinct motifs (green stacked bars) can be distinguished from U and D motifs (red and blue stacked bars, respectively). For each window, the TF motifs are shown in **Figures S2-S3** for mESCs and **S4-S6** for human WTC11 cells. In the U window, TFs that resemble U motifs are also detected, but the latter are less enriched in human compared to mouse cells (2.79% and 1.73% in WTC11 versus 25.5% and 11.85% in mEScs at accessible and inaccessible sites, respectively) as shown in **Figures S2A,** and **S4A-B**. Although CTCF is highly conserved between human and mouse, this result suggests that there are differences in their binding sites, such that in human there is less reliance on binding of ZFs 9-11 to the U motif. Motifs of other TFs enriched in the U and D windows and spacers and their preference for enrichment at accessible versus inaccessible sites are shown in **Figure 1B, S2-S3** for mESCs and **Figure 1B, S4-S6** for human WTC11 cells. The frequency with which each TF motif is found is shown as a percentage next to the motif in the figure. The majority of CTCF sites are enriched in one or more TF motifs, with each factor having a small contribution. It is pertinent that motifs in each window and group of TFs show similarities within and between TF families such as the ETS family in the U spacer, or the HOX family in the U window, indicating there is redundancy in the factors that could bind, thereby providing a potential mechanism for cell type-specific regulation. TF enrichment for each window in WT mESCs is shown in **Table S2A**, while in **Table S2B,** TF enrichment in WTC11 cells and other human tissues is shown to highlight that TF motif enrichment at CTCF sites is detected across different cell types, but as expected, motif enrichment of TFs varies, which could reflect each cell’s transcription program.

### The presence of TF motifs abutting CTCF binding sites is associated with stronger CTCF binding

To determine whether enrichment of TF motifs is associated with changes in CTCF signal strength, we compared the binding profiles of CTCF sites where we identified TF motifs at upstream or downstream regions to sites without TF motifs. The presence of TF motifs at accessible CTCF bound sites in the upstream and downstream regions, or both together were associated with increases in CTCF signals (**Figure 1C**). The overall CTCF background profiles were lower at inaccessible CTCF bound sites (**Figure 1C**) and the impact of TFs upstream or downstream had a distinct effect compared to accessible sites. The differential effect sizes could be explained by the predilection of TF classes found within specific windows as shown in **Figure 1B, S2-S3** and **S5-S6**. Importantly, enrichment of TF motifs is associated with a similar increase in CTCF binding strength in human WTC11 cells as observed in mESCs, suggesting that TFs play an equivalent role in stabilizing CTCF in the two species (**Figure 1D**). It is of note that motifs in up and downstream windows contribute in a distinct manner, in the two species, likely due to their distinct enrichment profiles.

In sum, the data in **Figure 1** shows that in human and mouse the majority of CTCF bound sites are enriched in one or more TF motifs. Their enrichment appears to be functionally important for increasing the stability of CTCF binding. Motifs within each window and group of TFs show similarities within and between TF families indicating redundancy in the cofactors that can bind. However, they are degenerate, thereby providing a mechanism for cell type-specific differences in regulation. These findings suggest that cell type-specific binding of CTCF might be influenced by a large pool of TFs that reflect the transcriptional program of each cell type, and our analyses provide a resource for identifying which TFs have the potential to influence CTCF binding and chromatin organization in different cell types (**Tables S2A, B**).

### Cobound TFs strengthen CTCF signals

We have shown that enrichment of TF motifs in the 35bps surrounding CTCF bound sites is associated with stronger CTCF signal (**Figure 1C** and **D**). To confirm the relationship between TF binding and CTCF signal strength, we analyzed ChIP-seq binding of individual TFs (KLF4, OCT4, MYC, SOX2 and YY1) in mESCs using data downloaded from O’Dwyer *et al*., and Laugwitz *et al*., ^20,21^. Our analysis, focused on CBSs with the consensus CTCF motif, highlighting sites at which CTCF binds alone, CTCF and TFs cobind and TFs bind alone. Heatmaps and profiles for accessible and inaccessible sites are shown in **Figure 2A** and **Figure S7**, respectively. The ChIP-seq profiles validate our motif enrichment analysis, demonstrating that individual TFs stabilize CTCF binding, with an increased effect at inaccessible sites for most TFs.

**Figure 2:**
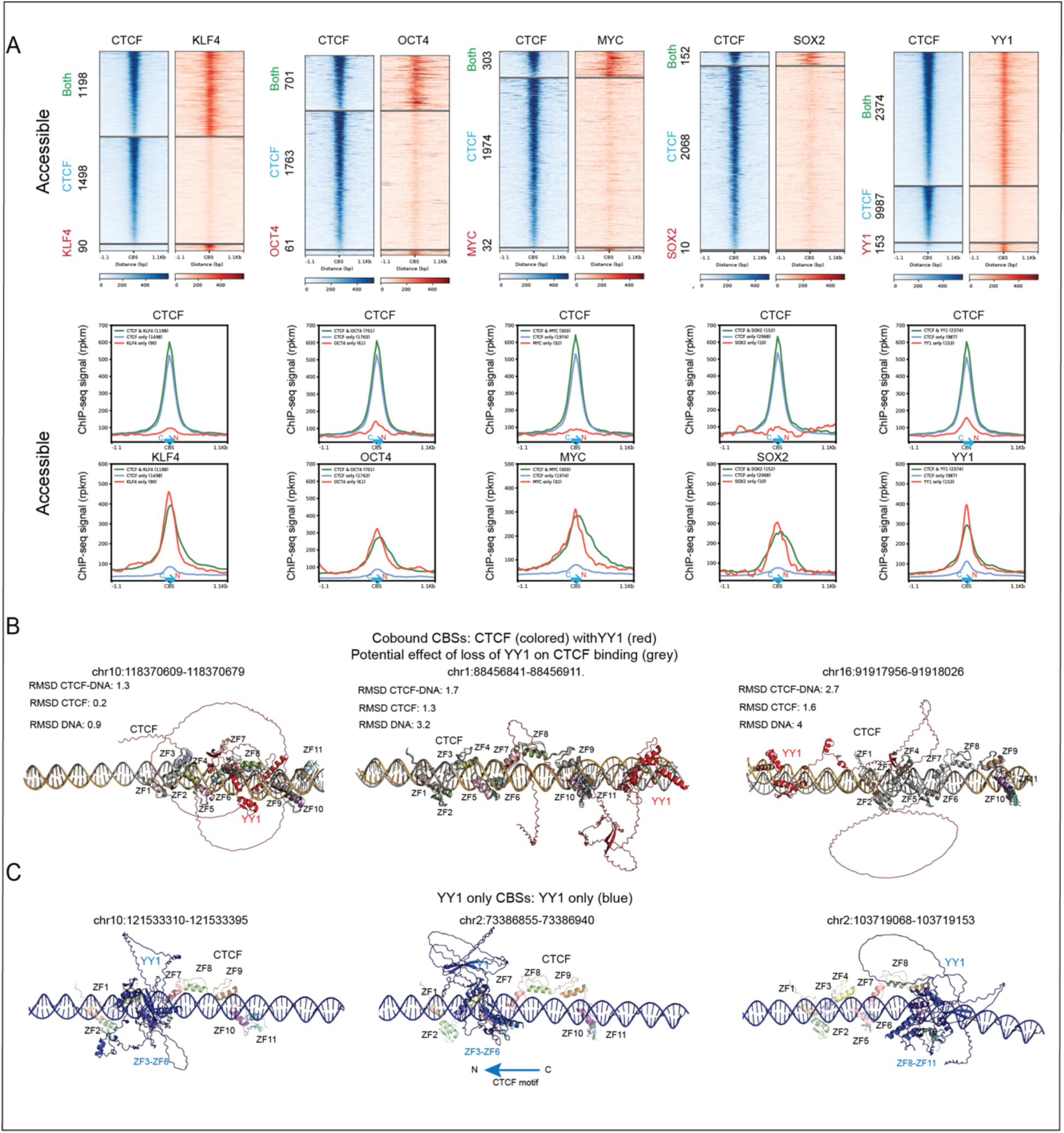
Individual TFs strengthen CTCF signal and disrupt CTCF-dependent nucleosome phasing. A: Heatmaps and profiles of CTCF (FLAG) and 5 TFs (including 4 pioneer TFs, KLF4, OCT4, MYC and SOX2) showing the effect of (i) TF binding on CTCF signal strength and (ii) CTCF binding on TF signal strength at sites where they are co-bound (green) compared to CTCF only (blue) and TF only (red) sites at accessible regions. This analysis is restricted to CTCF binding sites with the consensus CTCF motif and profiles are oriented on C to N terminal CTCF binding sites. The profiles at inaccessible sites are shown in **Figure S7**. **B**: AlphaFold3 predictions of the CTCF, YY1 and DNA complex at three cobound sites. The structures of CTCF with (colored) or without YY1 (grey) were aligned and compared by calculated the RMSD (root mean standard deviation) for the CTCF-DNA complex and separately for CTCF and DNA. **C**: AlphaFold3 predictions of the YY1 and DNA complex at three YY1 only bound CBSs. YY1 is shown in blue. CTCF binding predictions are overlayed to visualize the position of YY1 relative to CTCF. Predictions for the other TFs are shown in **Figure S8**.

The heatmaps and profiles also demonstrate that CTCF free CBSs occupied by TFs are associated with a stronger TF signal than at cobound sites for most TFs analyzed (**Figure 2A and S7**). The peak summits of the TFs are almost centered on the C to N terminal oriented CTCF motif, suggesting a competition between CTCF and the TF. At cobound sites however, we observe binding of the TFs at sites overlapping as well as on either side of the CBS.

### AlphaFold3 predicts CTCF/TF binding conformation at cobound and CTCF free CBSs

To get a clearer picture of how TFs and CTCF could affect each other’s binding, we turned to AlphaFold3 and analyzed TF binding at cobound and TF bound CTCF free CBSs for each of the five TFs, KLF4, OCT4, MYC, SOX2 and YY1. In **Figure 2B**, we show three examples of YY1 binding with CTCF at cobound sites. YY1 can bind in the Upstream spacer between ZFs 7 and 9 as well as the upstream or downstream regions. At these locations, there is little change in the CTCF conformation as reflected by the small change in the route mean square deviation (RMSD), however cobinding can affect DNA conformation which could result in changes in the CTCF-DNA interaction.

When we examined AlphaFold3 predictions of YY1 binding (blue) at CTCF free CBSs, we observed a direct competition between YY1 and CTCF at their predicted binding sites (**Figure 2C**), which could explain the absence of bound CTCF. Examples showing AlphaFold3 predictions of how CTCF and the remaining TFs could affect each other’s binding are shown in **Figure S8A-H**. As with YY1, each TF binds upstream, downstream or within the U spacer when cobound with CTCF, while at CTCF free CBSs, TFs tend to bind to sites overlapping the CBS, preventing CTCF binding. We also show examples of OCT4/SOX2 and MYC/MAX cobinding with CTCF. Interestingly, at the cobound locus shown in **Figure S8G,** the binding of SOX2 causes considerable DNA bending, which has been previously described ^22^. At this locus, the predicted binding is in the Upstream spacer between ZFs 7 and 9 which is flexible ^23^, allowing ZFs 9-11 to be properly positioned on the DNA despite the DNA bending, highlighting the importance of ZF8 as a spacer.

### Allele-specific analysis reveals both the cooperativity and competitiveness of CTCF-TF cobinding

To further validate the relationship between TFs and CTCF, we first downloaded the 145 ENCODE TF ChIP-seq sets available for the GM12878 cell line (**Table S3**) and found that 69% of the CTCF peaks overlap at least one TF, confirming that co-binding of CTCF with another TF is a frequent event. We then performed allele-specific analysis using ENCODE data from GM12878 to identify SNPs that impact CTCF or TF motifs (and therefore their binding affinity) (**Table S4**). In **Figure 3A, B** we show examples of heterozygous SNPs at two separate CTCF binding sites. In both cases CTCF binding is reduced on the allele with the base change, coupled with a loss of the cobound TF, overlapping RAD21 (a component of the cohesin complex) and to a lesser extent altered ATAC-seq signals. Thus, CTCF binding can affect binding of the TF, suggesting that in some cases CTCF binding might be required to open up the site for TF binding. Of note, the SNP in **Figure 3A** is linked in GWAS to colon cancer and cardiometabolic traits.

**Figure 3.**
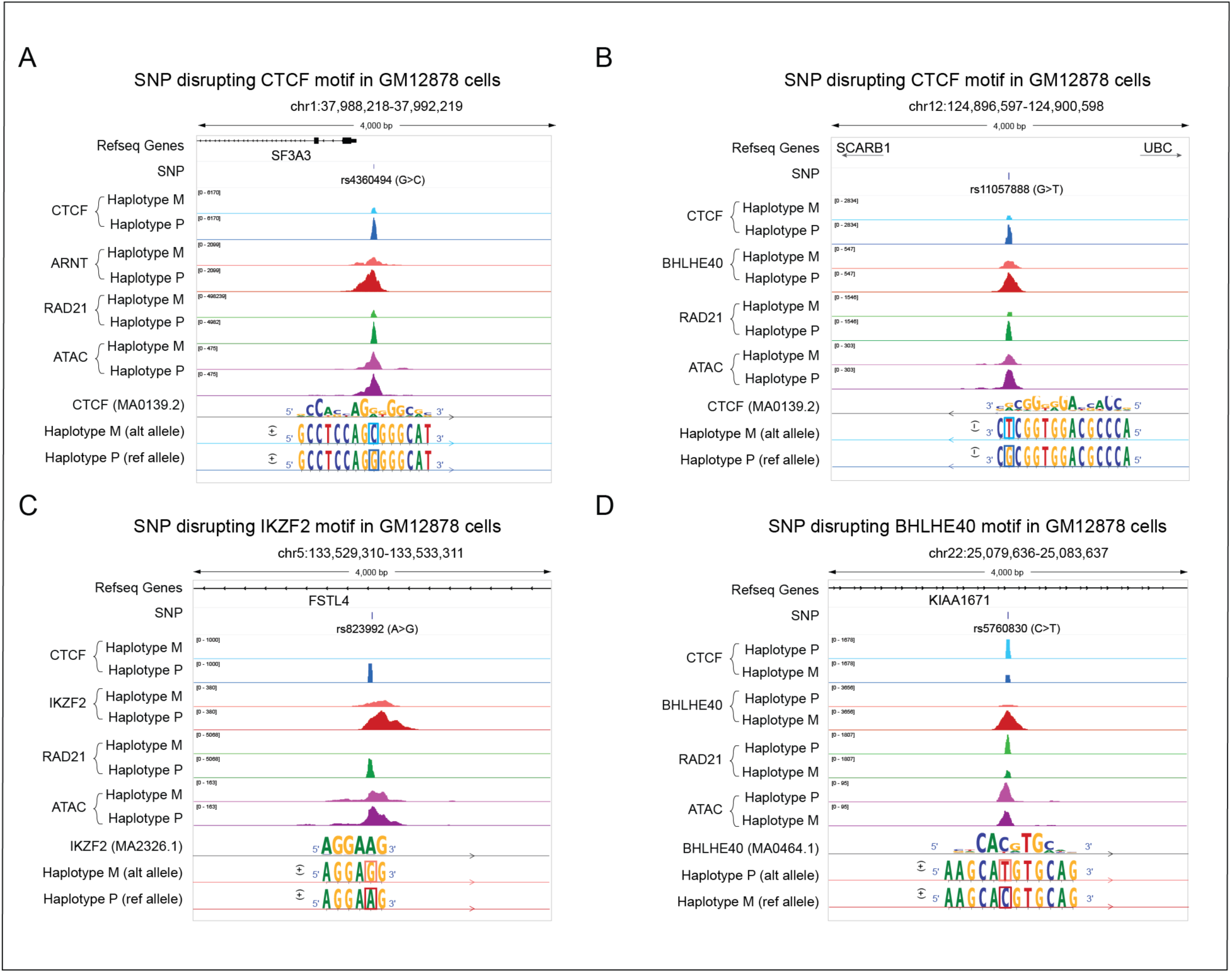
**SNPs reveal both the cooperativity and competitiveness of CTCF-TF cobinding**. **A**: Map showing allele-specific binding (ASB) of CTCF due to a SNP disrupting its motif on the maternal allele in the GM12878 cell line. RAD21 and ARNT binding are also disrupted on the CTCF unbound allele. The CTCF consensus motif and the sequence at this locus are shown on the bottom track along with the orientation. The rectangles highlight the heterozygous SNP. For visualization purpose, the sequence tracks are not to scale. M, maternal; P, paternal. **B**: Map showing ASB of CTCF due to a SNP disrupting its motif on the maternal allele, which is associated with the disruption of BHLHE40, cohesin binding and chromatin accessibility on the same allele. **C**: Map showing ASB of IKZF2 due to a SNP disrupting its motif on the maternal allele. CTCF, cohesin binding and to a lesser extent the chromatin accessibility are also disrupted on the IKZF2 unbound allele. **D**: Map showing ASB of BHLHE40 due to a SNP disrupting its motif on the paternal allele. At this locus, the unbound BHLHE40 allele is associated with CTCF and cohesin binding as well as more open chromatin. None of these regions overlap with known imprinting regions and the observed ASBs are likely be cis-dependent ASBs rather than parent-of-origin dependent.

Conversely, SNPs that affect binding of TFs on one allele can impact binding of CTCF at overlapping sites. In **Figure 3C**, the base change in the maternal haplotype impairs IKAROS binding, with a concomitant reduction in CTCF, RAD21 and ATAC-seq signals. Thus, in this setting binding of the TF is required for opening up the site to enable CTCF binding. In contrast, the SNP disrupting the BHLHE40 motif and binding on the paternal allele at the *KIAA1671* locus is permissive for CTCF binding and associated with concomitant binding of RAD21 and more open chromatin (**Figure 3D**). This highlights an example of a TF that binds competitively with CTCF and together with the data in **Figure 3B**, demonstrates that a single TF can have both a stabilizing and destabilizing effect as shown in **Figure 2A**.

Overall, we identified 131 regions with allele-specific binding of CTCF that are associated with allele-specific binding of a cofactor and where the presence of a SNP disrupting either the CTCF or the cofactor motif provides a causal explanation. In half of these cases, the perturbation introduced by the SNP indicates that TF co-binding impacts CTCF binding, while in the other half, the effect is reversed, confirming the reciprocal nature of TF-CTCF co-binding (**Table S4**). Additionally, we observed a destabilizing effect in 10% of the cases, suggesting that binding cooperativity is more prevalent than competitiveness.

### Transcriptional reprogramming contributes to cell state-specific CTCF binding

Our analyses demonstrate that cofactor binding can influence CTCF’s binding stability, however the impact of changes in transcriptional programs on CTCF binding have not been investigated. To address this question, we compared TF expression levels at accessible and inaccessible sites in human IPSCs. Overall, we found that TFs that bind motifs abutting CTCF were expressed at a higher level at accessible sites (**Figure 4A**) and that those sites are more sensitive to changes in TF expression levels compared to inaccessible sites, as shown by the significant difference in the correlation slopes in **Figure 4B**.

**Figure 4:**
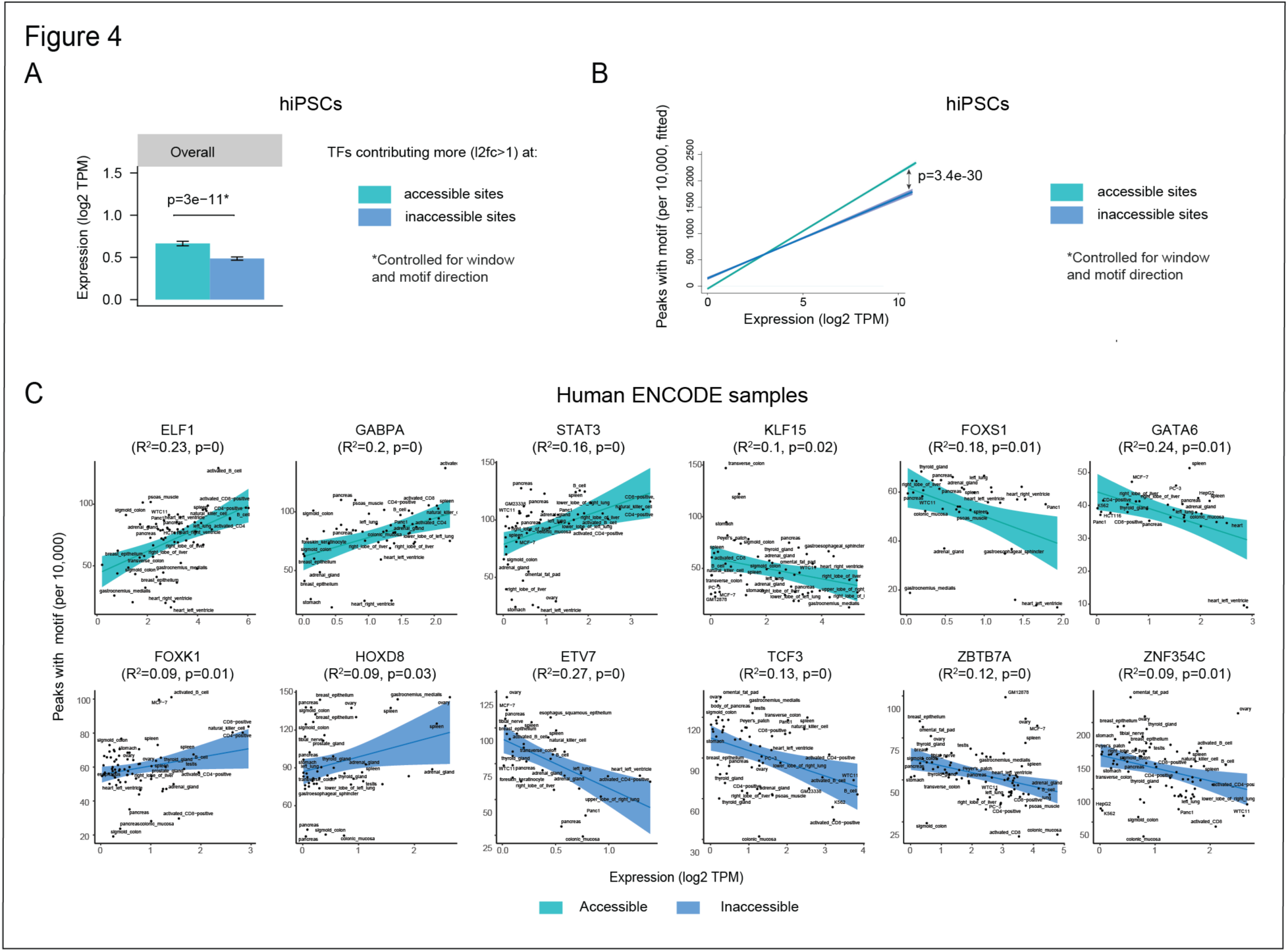
The percentage of TF motifs within 35 bp of CTCF consensus motif correlates with their expression level. A: Bar graph showing an increase in expression of TFs with a higher contribution at accessible versus inaccessible sites (≥2 fold differential contribution) in human iPSCs, after controlling for the differential effects of the gap window and direction relative to CTCF motif. The error bars correspond to the standard error computed using a multivariate linear model. **B**: Fitted correlation between overall TF expression and the percentage of CTCF sites found with motifs after controlling for the differential effect of windows and motif direction at accessible and inaccessible sites in human iPSCs. The shaded areas correspond to the 95%-confidence intervals. **C**: Examples of significant correlations between TF expression level and the percentage of CTCF peaks associated with the TF motif at accessible and inaccessible sites across human ENCODE cells and tissues. The shaded areas correspond to the 95%-confidence intervals.

To examine the effect of increasing transcriptional output on CTCF, we analyzed the effect of variations in TF levels across 84 different ENCODE cell lines on their enrichment at CTCF binding sites (**Table S2B**). We identified a subset of TFs whose expression levels showed a significant positive or negative correlation with motif enrichment around bound CTCF sites (**Figure 4C and Table S5**). This suggests that, in addition to context-specific effects, TFs might exert intrinsic stabilizing or destabilizing influences on CTCF binding and that the enrichment or depletion of a particular TF at CTCF sites in each cell type is dependent on its expression level, highlighting a mechanism by which cell type-specific transcriptional programs can impact CTCF binding profiles.

### CTCF mediates nucleosome repositioning around the CBS

To ask how CTCF binding affects nucleosome positioning in a context-specific manner, we performed long-read Nano-NOMe-seq in mESCs, which allowed us to assess nucleosome positioning using exogenous 5mC at HGC (GC excluding CGC) at the single molecule (SM) level. To assess the effect of CTCF motif orientation and to discriminate molecules with occupied and unoccupied CTCF binding sites, we restricted our analyses to CTCF peaks with GpC-containing CTCF motifs after confirming that nucleosome occupancy at these sites were similar (**Figure S9A**).

Unlike other similar analyses, we separated our profiles into occupied and unoccupied molecules. At CTCF ChIP-seq peaks, a substantial fraction of molecules was temporarily unoccupied, which is expected since DNA-protein interactions are dynamic. By comparing temporarily unoccupied to occupied molecules at CTCF bound sites - thereby controlling for context-specific confounding factors between bound and unbound CBSs - we confirmed previous findings demonstrating the displacement and phasing of flanking nucleosomes after CTCF binding (**Figure 5A**). Nucleosomes appeared less phased at inaccessible compared to accessible CTCF sites and exhibited slightly shorter periodicity as well as a reduced CTCF flanking nucleosome-free region (NFR) (**Figure 5A**).

**Figure 5:**
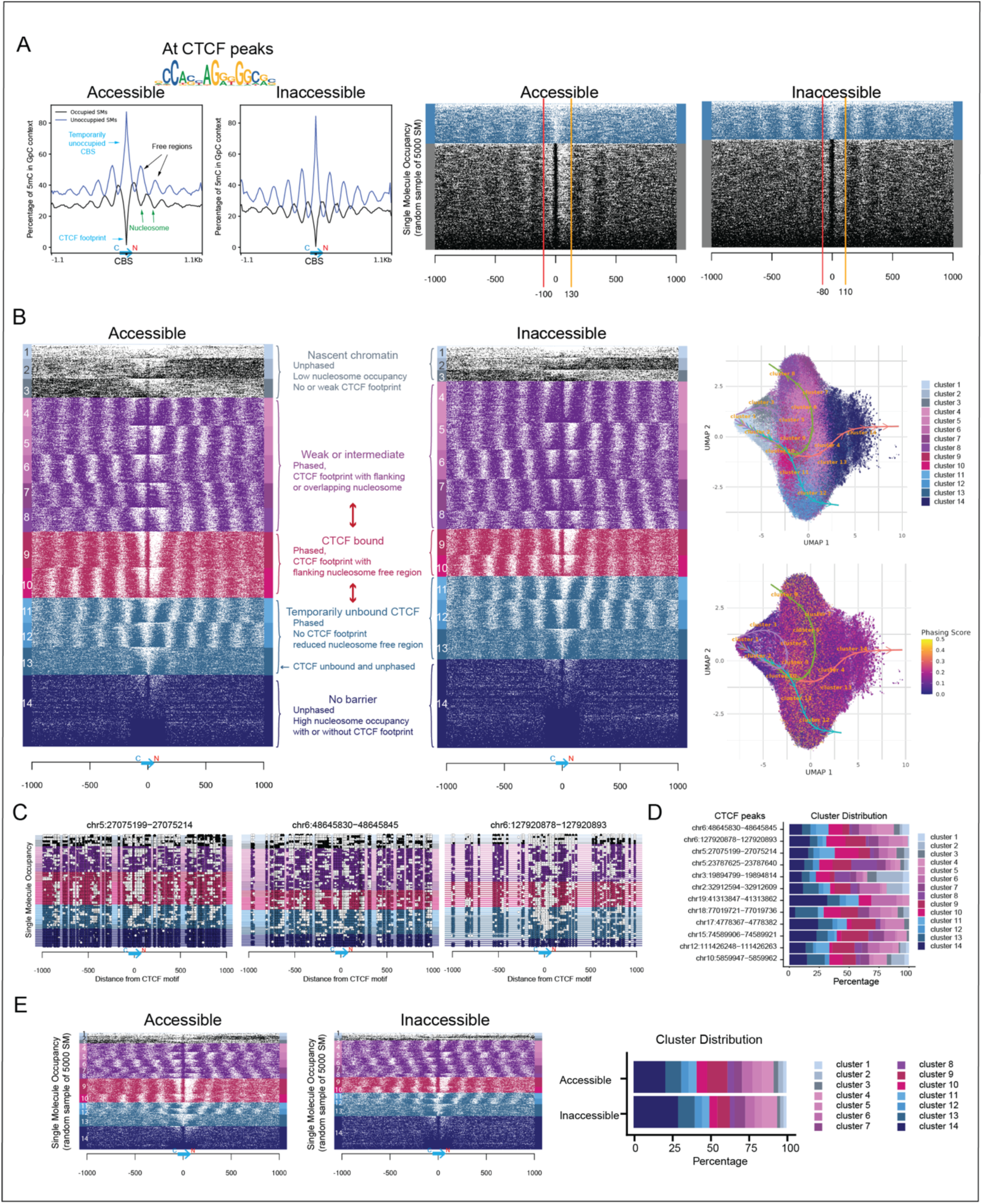
Subdivision of CTCF binding sites into 14 clusters reveals distinct phases of binding. A: Profiles and single molecule heatmaps showing the nucleosome occupancy (NO) and CTCF binding footprinting from nano-NOMe-seq for accessible and inaccessible CTCF sites. Both occupied (black) and temporarily occupied (blue) molecules at CTCF bound sites (ChIP-seq peaks) are shown. The data are centered on the 15-bp oriented consensus motif. The y-axis of the profiles shows the percentage of exogenous 5mC detected in an HGC context. High exogenous methylation reflects TF or nucleosome free chromatin accessible to the GpC methyltransferase, while lower methylation reflects occupied regions. The heatmaps show the nucleosome occupancy and CTCF footprinting at the single molecule (SM) level for a random sample of 5000 molecules. Black (or blue) and white dots represent occupied and free GpCs, respectively. The red and orange lines correspond to the stable position of the first upstream and downstream nucleosome flanking the CTCF motif. **B**: SM heatmaps of 5000 molecules per condition and genomic context showing the nucleosome occupancy and CTCF binding footprinting at accessible and inaccessible CTCF sites by cluster (left), and UMAP of the SMs colored by cluster and phasing scores (right). The arrowed lines show the SM trajectories identified by Slingshot. **C**: Example of heatmaps showing the nucleosome occupancy and CTCF binding footprinting at three individual CTCF peaks. **D**: Bar graph showing examples of cluster distribution at individual CTCF peaks. **E**: SM heatmaps and bar graphs showing the representative proportion of each cluster at accessible and inaccessible sites.

### Subdivision of CTCF binding sites into 14 clusters reveals a continuum of binding states at aggregated and individual peaks

To further understand the binding dynamics of CTCF binding on chromatin, we performed unsupervised clustering of the single molecules, incorporating data from all experimental conditions in mESC across both accessible and inaccessible regions. This generated 14 clusters, based on 50 bp-binned nucleosome occupancy within 1 kb of CBSs (**Figure 5B**). Differential enrichment of each cluster in specific conditions and genomic contexts underscores the biological relevance of each (**Figure S9B**). The UMAP projections of clustered SMs revealed that UMAP 1 correlates with global nucleosome occupancy whereas UMAP 2 partially captures differences in nucleosome phasing patterns, despite only subtle variations in phasing scores (**Figure 5B**, right and **S9C**).

Trajectory inference using Slingshot ^24^ helped define the relationships and dynamics among clusters. For this analysis, we set cluster 1 as the starting point, as the molecules in this cluster - along with clusters 2 and 3 (black clusters) - exhibit the characteristics of highly accessible, newly replicated chromatin, previously shown to lack well-defined nucleosome organization ^25^. Interestingly, cluster 1 displays a weak CTCF footprint with no surrounding nucleosome phasing, while clusters 2 and 3 exhibit weak phasing either on the left or right of the CTCF footprint, a distinction that could reflect the direction of replication. This observation suggests that CTCF can act as a barrier during redeposition of nucleosomes on nascent chromatin and might help to organize nucleosomes on newly replicated DNA.

Slingshot identified a bifurcation point corresponding to stable CTCF binding (red clusters 9-10) with two divergent branches (green and blue lines) along the UMAP 2 axis, leading to violet (clusters 4-8) and light blue (clusters 11-13) groups, respectively (**Figure 5B**). This branching pattern may reflect alternative temporary nucleosome configurations associated with CTCF binding. Indeed, the red clusters, enriched at strong CTCF peaks, show a pronounced CTCF footprint and well-defined flanking NFR, consistent with CTCF-associated nucleosome repositioning ^10^. The **violet** clusters (4-8) show a CTCF footprint on the left, middle or right side of a nucleosome, supporting findings that CTCF can bind its motifs regardless of nucleosome position ^26^. In these clusters, the close proximity of nucleosomes to the CTCF motif and the presence of partial NFRs suggest a less stable CTCF binding – supported by the enrichment in weak CTCF peaks - or intermediate states prior to, or during the action of nucleosome remodelers by CTCF ^10^. The **light blue** clusters (11-13) maintain nucleosomes in an organized, phased manner but with a narrower NFR around unbound CTCF sites (lacking a footprint), possibly reflecting temporally unbound CTCF. A third branch (orange) along UMAP1 leads to the **dark blue cluster** (14), which is enriched in the CTCF-depleted condition following IAA treatment. This might represent sites at which CTCF rebinds weakly or does not return at all, leading to disruption of nucleosome phasing. Finally, the purple path reflects the cyclical process between nascent and mature chromatin configurations.

Importantly when we focus on individual CTCF peaks we can detect all fourteen clusters, as shown for three representative examples (**Figure 5C**). These analyses are important for demonstrating that the clusters are not simply an aggregation of data from different binding sites enriched for one cluster or another across the genome. Instead, they represent a continuum of different states of CTCF binding at each peak. The proportion of each cluster is shown for twelve individual binding sites (**Figure 5D**).

Analysis of clustering at accessible and inaccessible sites highlights that the narrowing of the NFR surrounding CTCF seen in **Figure 5A**, is reflected by a decrease in the red clusters (strongly bound CTCF) and an increase in Cluster 14 (unbound or weakly bound CTCF with no nucleosome repositioning (**Figure 5E**). Additionally, a reduced ability to reposition nucleosomes is reflected by a more robust nucleosome presence on either side or overlapping the CBS (violet clusters). In sum, single molecule nano-NOMe-seq data reveals the dynamics of CTCF binding states at both aggregated and individual peak level, highlighting CTCF’s ability to reposition nucleosomes to create a NFR surrounding its binding sites and to act as a barrier that facilitates nucleosome phasing, both of which are impaired at inaccessible CBSs.

### Unoccupied CBSs are bound by nucleosomes or other factors

We next analyzed CBSs that are not bound by CTCF in mESCs. Most of the molecules at those sites are occupied (59% and 72% respectively at accessible and inaccessible sites), either by a nucleosome or a TF as suggested by the persistence of a narrow footprint (**Figure 6A**). Nucleosomes are still phased around the CBS but there is a noticeable increase in the proportion of violet and light blue clusters and a decrease in the red cluster compared to CTCF bound sites (**Figure 5E**). These data indicate that binding of TFs to CTCF free sites can still phase nucleosomes, but these are not associated with a NFR flanking the CBS. As before, we see a more robust nucleosome presence at inaccessible sites, consistent with less TF binding at these regions.

**Figure 6:**
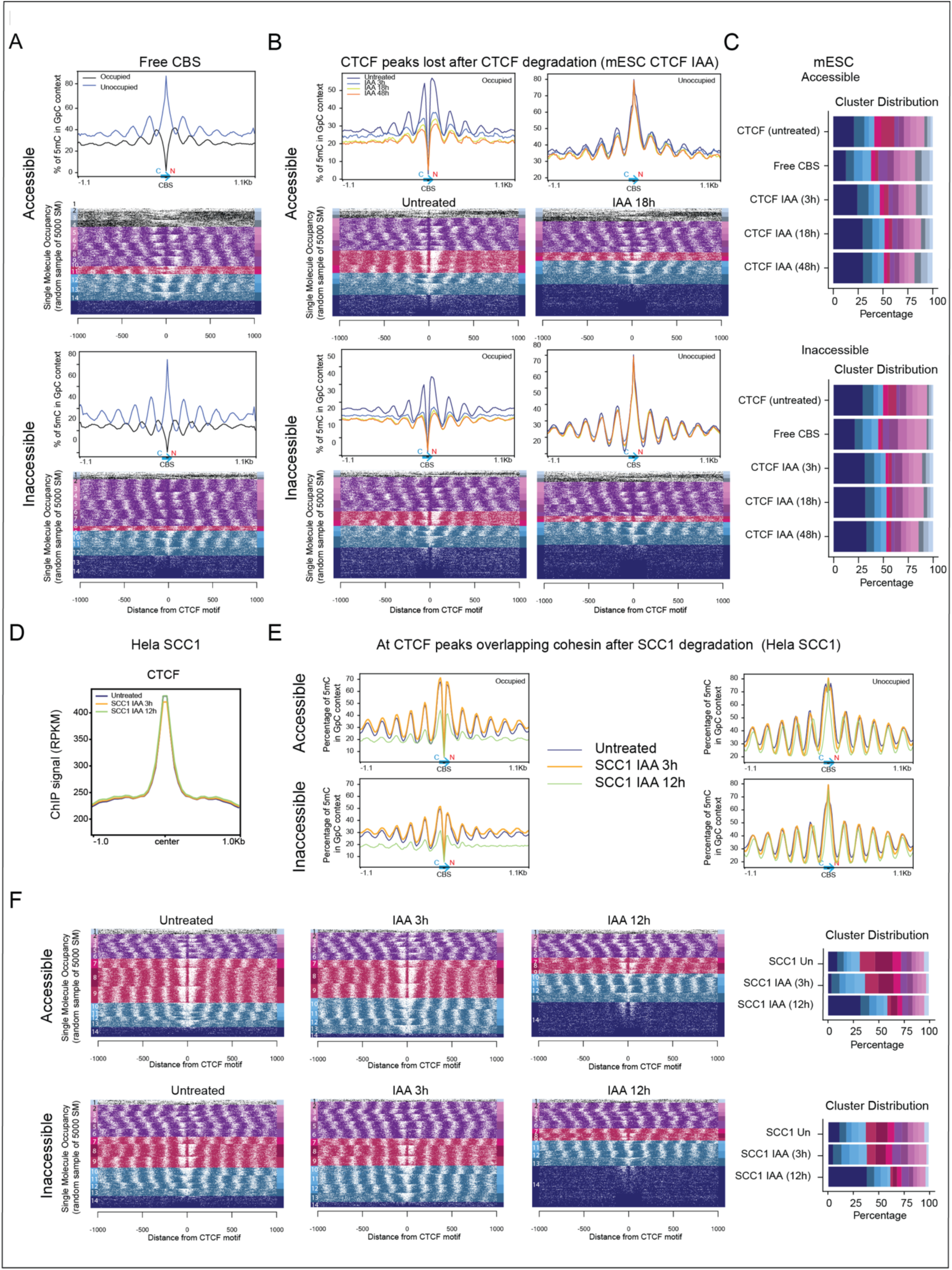
***SCC1 mediates CTCF associated nucleosome repositioning and phasing*. A:** Profiles and SM heatmaps stratified by cluster showing the nucleosome occupancy and CTCF binding footprinting for CTCF-free and motif-containing accessible and inaccessible CBSs. CTCF-free binding sites are defined as FIMO-detected CTCF motifs without CTCF or SMC3 ChIP-seq peaks. **B:** Profiles (for untreated, 3, 18 and 24 hours) and heatmaps of untreated and 18 hour IAA degraded mESCs at accessible and inaccessible sites. The heatmaps for 3 and 48 hours are shown in **Figure S10**. **C**: bar graphs showing the distribution of each cluster at CTCF peaks, CBS and after CTCF degradation (IAA). **D**: CTCF ChIP-seq profiles in untreated human Hela cells and after SCC1 degradation at 3 and 12 hours. **E:** Profiles of nucleosome occupancy in untreated human Hela cells and after SCC1 degradation (3 and 12 hours) at accessible and inaccessible sites. **F**: SM heatmaps and bar graphs showing the distribution of each cluster in untreated Hela cells and after IAA degradation.

### CTCF is required to reposition and phase nucleosomes

To confirm the effect of CTCF on nucleosome phasing and repositioning, we used an mESC CTCF auxin inducible degron ^9^. Loss of CTCF protein at 3, 18 and 48 hours resulted in an increasing reduction in nucleosome repositioning at CBSs (profiles in **Figure 6B**). The more granular analyses of the heatmaps and barplot show an increased proportion of cluster14 (dark blue), with a less marked footprint at the CBS over time (**Figure 6B, C** and **Figure S10A**). As expected, there is a notable decrease in the representation of the strong, red clustered CTCF binding sites and at the violet clusters, we detect a weaker CTCF footprint and a more robust nucleosome presence (**Figure S10A**). Interestingly, after CTCF degradation, CBSs show similar proportions of violet, red and light blue clusters to CTCF free sites (**Figure 4A**). Given these similarities, we speculate that the weak footprint at CTCF degraded sites could be due to the presence of incoming, or previously bound TFs, although we cannot rule out that some of this footprint is due to leftover, undegraded or weak CTCF ^27^.

### SCC1 mediates CTCF associated nucleosome repositioning and phasing

Since CTCF degradation leads to loss of both CTCF and cohesin at CBSs (**Figure S10B**), the CTCF degron experiment could not be used to determine which factor is responsible for mediating the changes we observed after CTCF degradation. To address this question, we used an auxin inducible degron of SCC1 (a component of the cohesin complex) in the human Hela cell line ^28^. Since cohesin is required for mitosis, we assessed early degradation time points, 3 hours and 12 hours, where we could still collect live cells. Alongside nano-NOMe-seq we performed CTCF ChIP-seq. CTCF binding profiles remained the same as untreated cells after IAA treatment at 3 hours and 12 hours (**Figure 4D**). In contrast, the nucleosome profiles for occupied and unoccupied sites show that although there is little change in nucleosome clustering at the 3-hour time point, despite unchanged CTCF profiles, we could detect a reduction in nucleosome repositioning and phasing at CBSs by 12 hours (**Figure 4E**), similar to that observed after CTCF degradation (**Figure 4B)**. Looking at the heatmaps and barplots (**Figure 4F)** at both accessible and inaccessible sites there is a noteable reduction in the strongly bound CTCF fraction (red clusters) and an increase at weakly bound CTCF sites characterized by disorganized nucleosomes surrounding the CBS (dark blue clusters). Violet clusters are also reduced but more so at inaccessible regions, and at these sites there is a stronger nucleosome presence surrounding the CBS, obscuring the CTCF footprint in most places. That the 12-hour time point effect occurs in the background of unaltered CTCF binding, indicates that cohesin plays a dominant role in nucleosome repositioning and phasing. However, CTCF alone may exert an additional weaker effect, as the red clusters are not reduced to the same extent as when CTCF is depleted and there is a persistent NFR in the dark blue cluster. We speculate that the delay in loss of nucleosome repositioning at 3 hours could be because nucleosome repositioning needs to be reset during S phase ^29^.

In sum, we found that CTCF free CBS, CTCF and SCC1 degradation all give rise to similar clustering profiles with a decreasing NFR over time, indicating that a reduction or absence of CTCF binding can allow binding of TFs that can phase, but are unable to reposition surrounding nucleosomes. Thus, cohesin and loop extrusion could mediate nucleosome positioning and phasing by creating or stabilizing barrier effects at CTCF anchors.

### CTCF signal strength contributes more to nucleosome repositioning and insulation than accessibility

When we compare the clustering of strong versus weak CTCF signals and inaccessible versus accessible sites, we observed that signal strength gives rise to a bigger difference in the proportion of red clusters (strong CTCF binding in a NFR), compared to accessibility (**Figure 7A**). Moreover, the differential effects of CTCF signal strength were similar at both accessible and inaccessible sites, suggesting that CTCF signal strength contributes to nucleosome repositioning independent of accessibility. We previously performed a stratified analysis showing that CTCF signal strength, which was associated with cohesin stalling, also contributes more to insulation than accessibility ^9^. Here we highlight that a reduction in nucleosome repositioning at weak versus strong sites is associated with a corresponding reduction in chromatin insulation compared to that at inaccessible versus accessible sites (**Figure 7B**).

**Figure 7:**
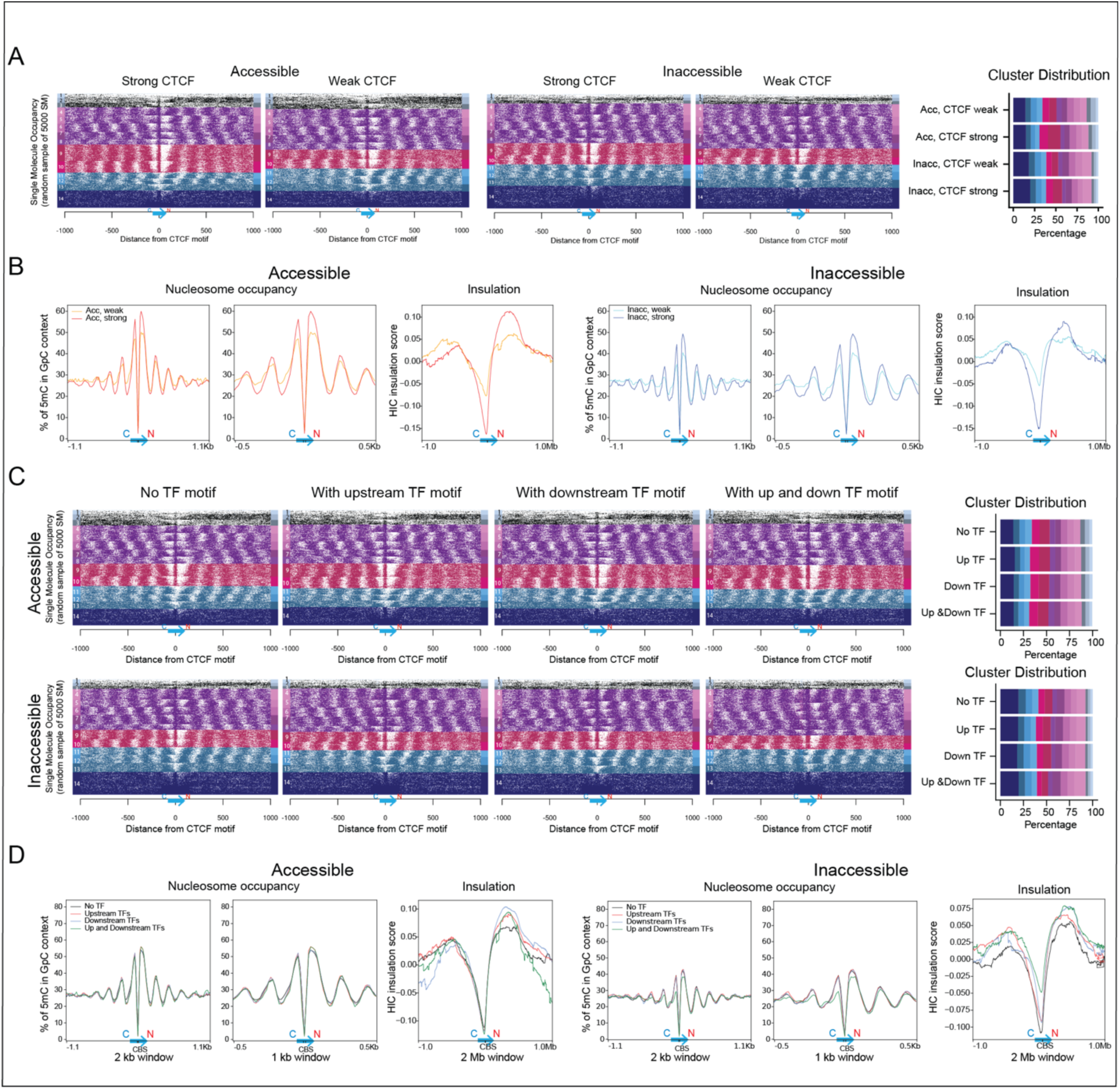
The CTCF flanking NFR is associated with insulation. **A.** Clustered heatmaps of SMs stratified by chromatin accessibility and CTCF signal strength. **B:** Profiles comparing the nucleosome occupancy and insulation score between weak versus strong CTCF sites stratified by chromatin accessibility. The averaged insulation score generated at 10 kb resolution from HIC data is shown for each genomic context. **C**: Heatmaps of SMs stratified by clusters at accessible and inaccessible CTCF peaks in the absence or presence of TF motifs. The bar graphs show the distribution of each cluster by genomic context. **D**: Profiles comparing the nucleosome occupancy and insulation score between CTCF-alone and cobound sites stratified by chromatin accessibility.

### Surrounding motifs affect CTCF-mediated nucleosome repositioning and insulation independent of signal strength

The presence of TF motifs are associated with differential effects on CTCF binding (**Figure 1C**). Although CTCF signal strength is increased at accessible sites, we observe a minimal change in nucleosome repositioning and chromatin insulation (**Figure 7C, D**). The same is true at inaccessible sites, except when TF motifs are enriched on either side of CTCF. Despite a slight increase in CTCF signal strength at these sites, both nucleosome repositioning and chromatin insulation are reduced. Together, these data indicate that CTCF and cohesin signal strength can be uncoupled from the size of the NFR surrounding CTCF, while the association between nucleosome repositioning and insulation is maintained. This finding highlights that nucleosome repositioning is a more reliable predictor of chromatin insulation than CTCF signal strength. The data also demonstrate that TF binding can interfere with nucleosome repositioning and chromatin insulation and we speculate that the presence of TFs could be a way to fine tune the effect of CTCF signal strength on these two processes.

## Discussion

Although it is known that a subset of CBSs is altered in a state-specific manner, the mechanisms underlying this control have not been fully elucidated. Here we used CTCF ChIP-seq in mESCs combined with CTCF ChIP-seq from ENCODE data in a wide range of human cell types to (i) characterize accessible versus inaccessible sites, and (ii) determine what factors influence CTCF binding stability and function within these two chromatin environments. Although CTCF is highly conserved in mice and humans, we found different features associated with their respective binding sites. Specifically, mouse cells have proportionately more inaccessible binding sites compared to human cells. In parallel, TF motifs are enriched at inaccessible sites in mouse, which we have previously shown generally have weaker CTCF signals and reduced cohesin overlap compared to accessible sites ^9^. Since TF cobinding stabilizes CTCF, we speculate that their enrichment at inaccessible sites in mouse, restores the balance of strong versus weak CTCF/cohesin signals in the two species, compensating for the relative reduction of accessible sites in mouse.

CTCF has been shown by us and others to act as a pioneer factor, binding inaccessible as well as accessible chromatin ^9,30^. To follow up on previously published studies hinting at a role for TFs in cell state specific binding of CTCF, we performed a comprehensive analysis of enrichment of TF motifs in the 35bp region of the CBS in mESCs as well as 85 ENCODE cell lines. We identified a wide range of motifs corresponding to expressed cell state-specific factors that could perform this function. In human and mouse, the majority of CTCF bound sites are enriched in motifs located within 35 bps of CTCF. Within a given window, enriched TF motifs show a strong similarity, suggesting redundancy in the regulation of CTCF binding. Furthermore, motifs are degenerate which could allow for differential effects of TFs from the same class or motif cluster. Given the redundancy between factors that can bind, we speculate that at a single CTCF binding site, the presence of distinct TFs in different cell types could fine-tune CTCF’s stability and its ability to block cohesin in a lineage or state specific manner. Indeed, we demonstrate that CTCF signals are strengthened at sites enriched with TF motifs and by analyzing the effect of a handful of cofactor TFs on CTCF binding profiles, we validate their stabilizing effect.

The significance of these findings is underscored by our analysis of human tissues from ENCODE, all of which have unique transcriptional programs with individual TFs expressed at different levels. Importantly, increases in the expression level of a TF can both increase or decrease the enrichment of its motif at CTCF bound sites, depending on whether the TF has a stabilizing or destabilizing effect. The relationship between TFs and CTCF were validated by allele specific analysis which demonstrate that the two factors have a reciprocal effect on each other’s binding such that SNPs affecting either the binding of CTCF or the TF will impact enrichment of the other factor, altering cohesin overlap and accessibility. SNPs in CBSs or TF motifs highlight both the cooperativity and competitiveness of CTCF-TF cobinding. Our finding that cell type-specific TFs can have a negative effect on CTCF binding addresses another long-standing question about why many accessible CBSs remain unbound. Here we show that binding of other factors can compete with CTCF to prevent its occupation.

AlphaFold3 predictions demonstrate that TF binding to CTCF free CBSs, overlaps the CBS, providing an explanation for their inability to cobind. In contrast, at cobound sites, binding of the TF occurs in the upstream or downstream regions, sites that do not interfere with CTCF’s occupation. Although AlphaFold3 cannot predict the impact of cobinding on CTCF stability, our ChIP-seq analyses confirm that this leads to a stronger CTCF signal, despite TF-mediated changes in DNA torsion and bending. However, binding of cofactors can alter nucleosome phasing, and we speculate that changes in DNA torsion and bending could be responsible for disrupting repositioning, and the outcome could be TF specific as previously shown ^22^. Cobinding of YY1 and CTCF occur at a subset of PAX5 binding sites in the *immunoglobulin heavy chain (Igh)* locus ^31^. This is of interest, because we previously showed that both PAX5 ^32,33^ and YY1 ^34^ are important for mediating chromatin looping and locus contraction in early B cell development to induce long range V-to-DJ rearrangements which is important for the generation of antibody diversity. In the absence of either factor, locus contraction and rearrangement are impaired, consistent with the idea that cofactors are important for CTCF mediated function.

Use of long read, nano-NOME-seq single molecule analysis allowed us to interrogate the impact of CTCF binding on nucleosome repositioning at a resolution other assays such as MNase could not achieve. By subsetting the data, we were able to show a continuum of CTCF binding states, starting with binding to highly accessible chromatin devoid of nucleosomes, which has been shown to reflect newly replicated chromatin ^25^. Binding of CTCF occurs early and helps to organize nucleosomes around the binding site leading to ordered nucleosome phasing and repositioning, which likely requires recruitment of chromatin remodellers, such as members of the mSWI/SNF family ^35^. The clustering shows that CTCF can bind with either its N or C terminal abutting a nucleosome or less frequently, overlapping a nucleosome. This is in contrast to the findings of Pugacheva *et al*., who posit that CTCF binds only at N terminal nucleosome entry sites ^35^.

At CTCF free sites, CBSs are occupied either by a nucleosome or TFs that can phase nucleosomes, but cannot reposition them. The effect of the TF on nucleosomes is more dynamic as reflected by less stable nucleosome positioning around CTCF, in line with the low residence time of of these factors compared to CTCF. Furthermore, there is a more robust nucleosome presence at inaccessible sites, consistent with less TF binding at these regions. Interestingly, we detect similar overall binding states when we degrade CTCF, suggesting that in the absence of CTCF, TFs can come in and bind at CBSs. This data further highlights that CTCF and TFs can bind competitively at CBSs, when their binding sites directly overlap.

While CTCF has been shown to act as a barrier that promotes nucleosome phasing and positioning at its binding sites ^36^, SCC1 depletion data suggests that cohesin plays a critical and dominant role in nucleosome repositioning and phasing at CTCF sites. Notably, the loss of phasing and repositioning observed following SCC1 degradation - despite unchanged CTCF binding strength - closely recapitulates the nucleosome configuration seen upon combined loss of CTCF and SMC3 binding at CBSs after CTCF depletion. This finding suggests that cohesin and loop extrusion might be responsible for the local nucleosome organization observed at CTCF anchors.

Clustering demonstrates that the proportion of chromatin with repositioned nucleosomes depends on accessibility as well as CTCF signal strength, both of which likely influence the recruitment of nucleosome remodellers. However our studies highlight that signal strength has a greater impact on nucleosome repositioning, and this is also reflected in a bigger impact on chromatin insulation than accessibility, confirming our previous findings ^9^. However, we also show that CTCF signal strength and nucleosome repositioning can be uncoupled in the presence of TFs, while the association between nucleosome repositioning and insulation is still maintained. Thus, nucleosome repositioning is a more reliable predictor of insulation than CTCF signal strength and we propose that use of single molecule nucleosome occupancy could be a powerful tool to predict effects on insulation.

Here we have shown that enrichment of a wide range of TF motifs stabilize CTCF binding and confirmed this by analyzing the ChIP-seq of a handful of factors at CBSs. Together, these studies define a mechanism by which transcriptional reprogramming during development and in disease specific states can determine cell type-specific CTCF profiles, linked to local and long-range chromatin organization.

## METHODS

### Cell line generation and culture

The construction of the vector for cloning transgenic, doxycycline-inducible expression of WT and mutant mouse *Ctcf* cDNA and gene targeting in mouse embryonic stem cells E14Tg2a (karyotype 19, XY; 129/Ola isogenic background) were performed as described in Do *et al.,* ^9^. Mouse embryonic stem cells and all clones derived from these were cultured under feeder-free conditions in 0.1% gelatin (Sigma ES-006-B) coated dishes (Falcon, 353003) as described in Do *et al.,* ^9^. Briefly, the cells were grown in DMEM (Thermo Fisher, 11965-118) supplemented with 15% fetal bovine serum (Thermo Fisher, SH30071.03), 100 U/ml penicillin - 100 μg/ml streptomycin (Sigma, P4458), 1 X GlutaMax supplement (Thermo Fisher, 35050-061), 1 mM sodium pyruvate (Thermo Fisher, 11360-070), 1 X MEM non-essential amino-acids (Thermo Fisher, 11140-50), 50 μM b- mercaptoethanol (Sigma, 38171), 10^4^ U/ml leukemia inhibitory factor (Millipore, ESG1107), 3 μM CHIR99021 (Sigma, SML1046) and 1 μM MEK inhibitor PD0325901 (Sigma, PZ0162).

SCC1-mEGFP-AID HeLa cell line in which SCC1 is tagged with an auxin induced degron was previously generated by Wutz et al.,^28^ and was kindly provided by Dr. Peters. The cells were grown in DMEM supplemented with 10% DBS (Thermo Fisher, 10371029), 0.2 M glutamine (Thermo Fisher, 25030081), penicillin/streptomycin and 1X non-essential amino acids (Thermo Fisher, 11140050)

### Induction of auxin inducible degradation of CTCF, SCC1 and doxycycline induced expression

For degradation of endogenous CTCF in mESCs, the auxin-inducible degron was induced by adding 500 μM indole-3-acetic acid (IAA, the chemical analog of auxin) (Sigma, I5148) to the media. Expression of WT CTCF transgene was achieved by the addition of doxycycline (Dox, 1 μg/ml) (Sigma, D9891) to the media. The cells were treated with IAA and/or Dox for 3h, 18h or 2 days. In Hela, SCC1 were degraded by addition of 500 μM IAA for 3h or 12h.

### Nano-NOMe-seq sample and library preparation

#### Nuclei extractraction, GpC methyl-transferase treatment and DNA extraction

After 2 days of IAA and Dox treatment on WT mESCs, 1 million cells were collected and washed with cold PBS at 300g at 4°C. As described in Battaglia *et al*., ^37^, nuclei were extracted using cold lysis buffer (10 mM Tris-HCl pH 7.4, 10 mM NaCl, 0.5 mM spermidine, 1.5 mM MgCl2, 0.1 mM EDTA, 0.25% IGEPAL CA-630) for 3-6 minutes after checking nuclei under a light microscope. The sample was centrifuged for 5 minutes at 800x g at 4°C, washed with 1 ml ice-cold nuclei wash buffer (10 mM Tris-HCl pH 7.4, 50 mM NaCl, 0.5 mM spermidine, 1.5 mM MgCl2, 0.1 mM EDTA), and centrifuged again for 5 minutes at 800x g at 4°C. The nuclei were resuspended in 564 µL 1X GpC MTase reaction buffer (NEB, B0227) and equilibrated to 37°C. This was followed by 6 minutes of enzyme treatment (300 µL 1 M sucrose, 34 µL 10X GpC methyltransferase reaction buffer, 3 µL 32 mM S-Adenosylmethionine (SAM) (NEB, B9003), 100 µL GpC methyltransferase (M.CviPI) (NEB, M0227L)), followed by another 6 minutes of enzyme treatment (50 µL GpC methyltransferase (M.CviPI), 3 µL 32 mM S-Adenosylmethionine (SAM)). An equal volume of pre-warmed 2X stop buffer (20 mM Tris-HCl pH 7.4, 600 mM NaCl, 10 mM EDTA, 1% SDS) was added and mixed well. Proteinase K (200 µg/mL) was added to the GpC methylation reaction and incubated for 4 hours at 56°C, followed by RNase A (100 µg/mL) for 30 minutes at 37°C, followed by an additional round of proteinase K (200 µg/mL) for 30 minutes at 37°C.

HMW DNA was extracted using phenol-chloroform and precipitated by isopropanol. Briefly, an equal volume of phenol-chloroform was added to the samples and mixed vigorously. The samples were centrifuged at 13,000 rpm for 10 minutes at room temperature. The aqueous phase was transferred to a clean tube, and 0.7-1 times the volume of isopropanol was added, mixed well, and incubated at room temperature for 30 minutes. The samples were then centrifuged at 13,000 rpm for 10 minutes at 4°C. The DNA pellet was washed with 1 ml pre-chilled 70% ethanol. After removing all the ethanol, the DNA was dissolved in EB buffer and incubated at 37°C for 30 minutes. The final DNA concentration was quantified using Nanodrop 2000, and molecular weight was measured by genomic DNA screentape (Agilent, 5067-5365).

#### Nano-NOMe-seq library preparation

For Nano-NOMe-seq, the DNA fragment was sheared to an average size of 20kb with the Covaris G-Tube (#520079) in an Eppendorf 5415D tabletop microcentrifuge following the manufacture’s protocol. Sample size was confirmed on genomic screentape with the Agilent Tapestation system. The sheared DNA (1.5ug) was then input into the SQK-LSK114 library prep from Oxford

Nanopore to create the Nanopore library. Components were used from the LSK114 kit along with several buffers and enzymes from New England Biolab (NEB) following the LSK114 protocol from Oxford Nanopore. Following library prep, the sample concentration was verified on the Qubit system using the dsDNA High Sensitivity Kit. The concentration and basepair size were used to calculate Femtomolar concentration for loading. The sample was resuspended with library loading beads and loaded at 150-200 femtomolar concentration onto the Promethion R10.4.1 flow cell according to the LSK protocol. Sample were sequenced on the R10.4.1 Promethion flow cell using the Oxford Nanopore Promethion P24 system with a sequencing run time of 72 hrs. Standard settings were applied during the run setup, special settings were applied and fragments less than 1kb rejected and methylation detection was set for both 5mC and 5hmC along with the super accurate basecalling feature turned on.

#### ChIpmentation

ChIPmentation was performed in the Hela cell lines as described in Do *et al.* ^9^. Briefly, 10 millions cells were double cross linked using 25mM EGS (ethylene glycol bis(succinimidyl succinate); Thermofisher #21565), followed by addition of 1% formaldehyde (Tousimis #1008A). Chromatin was sheared using a bioruptor (Diagenode) for 15 minutes. Mouse anti-CTCF (Active Motif, 61311) and Rabbit IgG (abcam, ab37415) antibodies were used for immunoprecipitation. Tagmentation were performed using 1 μl Tagment DNA Enzyme from the Tagment DNA Enzyme and Buffer Kit (Illumina #20034198). Purified DNA (20 μl) was amplified and barcoded using indexed and non-indexed primers and NEBNext High-Fidelity 2X PCR Master Mix (NEB M0541). DNA was purified using Agencourt AMPure XP beads (Beckman, A63881) to remove fragments larger than 700 bp as well as the primer dimers. Library quality and quantity were estimated using Tapestation (Agilent High Sensitivity D1000 ScreenTape #5067-5584 and High Sensitivity D1000 reagents #5067-5585) and quantified by Qubit (Life Technologies Qubit™ 1X dsDNA High Sensitivity (HS) #Q33230). Libraries were then sequenced with the Novaseq6000 Illumina technology according to the standard protocols and with around 200 million 150bp paired-end total per sample.

### Datasets and Data Processing

#### ChIP-seq, RNA-seq and ATAC-seq dataset

The CTCF, SMC3 ChIP-seq, RNA-seq and ATAC-seq data for WT mESCs were previously generated and processed as described in Do *et al.*, ^9^ using the Seq-N-Slide pipeline ^38^. Hela ChIPmentation data were processed using the same pipeline.

For ENCODE human samples, processed data for samples having CTCF ChIP-seq, RNA-seq and ATAC-seq performed on the same biosample were downloaded from ENCODE. The accession numbers of the dataset used in this work are listed in **Table S1A, S1B, S3**. CTCF peaks were scanned for CTCF consensus motif (JASPAR MA0139.2) ^39^ using FIMO ^40^ with a p-value cutoff of 5e10^-5^. Only samples with at least 20% of motif containing CTCF peaks were kept for downstream analyses. Accessible CTCF peaks were identified by overlapping CTCF with ATAC peaks using bedtools ^41^.

For the GM12878 cell line, we downloaded an additional set of 145 ENCODE TF ChIP-seq narrow peak files to assess the percentage of CTCF binding sites with TF co-binding (the accession numbers are listed in **Table S3**. To analyze allele-specific binding the BAM files for CTCF, RAD21, TFs and ATAC-seq were also downloaded (the accession numbers are listed in **Table S3**). The phased genotyping data of GM12878 was retrieved from the Platinium Genomes project ^42^ (accession number PRJEB3246).

To validate the effect of TF co-binding, we downloaded KLF4, OCT4, SOX2, MYC, YY1 and input ChiP-seq data (fastq files) generated in mESCs from GEO ^20^ ^21^. The data were processed using the Seq-N-Slide pipeline ^38^ as described in Do *et al*., ^9^. Briefly, the reads were aligned to mm10 genome with Bowtie2 ^43^ and narrow peaks were called using MACS2 ^44^ in pair-end mode and with the input as control.

The CTCF-oriented heatmaps and profiles were generated using the computeMatrix scale-regions function from deeptools ^45^ using a 15 bp bin and a region body length of 15 bp which corresponds to the length of CTCF core motif.

#### Hi-C data

The HIC data were previously generated using Arima HiC kit (A510008) and processed using HiC-Pro and HiCExplorer as described in Do *et al*., ^9^. The HIC matrices were generated at a 10 kb resolution. The insulation scores were computed using GENOVA ^46^ and the CTCF-oriented profiles generated using deeptools ^45^ with a 1 kb bin and a region body length.

#### Nano-NOMe-seq processing

Base modification calling were performed on pod5 files using dorado basecaller ^47^ using the “-- min-qscore 10 sup,5mC_5hmC” option, followed by alignment using dorado aligner against the mm10 reference genome. Reads with less than 10 MAPQ score were filtered out using samtools^48^. Methylation in HGC context was called using dorado pileup using the “--combine-strands” option. Single molecule (SM) methylation was extracted using modkit extract with the “--read-calls-path” option ^49^. GpC overlapping the ENCODE blacklisted regions ^50^ were excluded as well as sites with more than 100x coverage.

CTCF-oriented GpC methylation profiles were generated using deeptools and single molecule heatmaps generated using R. Single molecules with occupied or unoccupied CTCF were defined as molecule containing a unmethylated or methylated GpC in the motif, respectively. No bias was observed in the nucleosome positioning of molecules with and without GpC containing motifs (**Figure S4A**). SM Heatmaps were created in R at the single molecule and GpC level using the GpC methylation status called by modkit extract after centering the reads on the middle of CTCF consensus motif.

### Downstream Analyses

#### Motif logos and Motif spacing enrichment analysis

Motif logos of the aggregated sequences centered on CTCF motif detected by FIMO ^40^ were created using the R package, ggseqlogo ^51^. Significantly enriched spacings of TF motifs in the vicinity of CTCF bound motifs were identified using SpaMo ^52^. The motif positional weight matrices (PWM) were downloaded from JASPAR 2024 (CORE vertebrates non-redundant dataset) ^39^. The PWM for the U and D motifs were derived from Nakahashi *et al.,* ^17^. The analyses were performed using the 500 bp window centered on each bound CTCF motif stratified by chromatin accessibility. TF motif clustering and trees were performed using the R package, universalmotif ^53^ for each window defined as follow: upstream spacer, U window and further upstream, 0 to 5bp, 6-8 bp and more than 9bp upstream, respectively; downstream spacer, D window and further downstream, 0 to 9 bp, 10-12 bp and more than 13 bp downstream respectively.

#### Association between TF motif enrichment and TF expression

To ask whether TFs more enriched at accessible sites compared to inaccessible sites were more expressed in human iPSCs, accessible enriched TFs were defined as TFs with a 2-fold increase in the percentage of motif found within 35 bp of the CTCF motif identified by the SpaMo analysis. A multivariate linear regression using a backward-stepwise selection was then performed to test the association between TF expression level (log2 TPM) and chromatin accessibility controlled for the epistatic effect of the motif orientation and window of enrichment (relative to the CTCF motif). The adjusted expression level by accessibility was estimated by computing the model margins and the p-value by performing the pairwise comparisons of predictive margins using STATA v17.

To test the overall correlation between TF expression and motif enrichment, we performed an aggregated analysis across ENCODE human samples (**Table S1A**). The expression level of TFs and percentage of motif were summed by sample, accessibility, window of enrichment and motif orientation. A multivariate linear regression using a backward-stepwise selection was then performed to test the correlation between the aggregated percentage of enriched motifs and TF expression level (log2 TPM) controlled for the epistatic effect of chromatin accessibility, the motif orientation and window of enrichment (relative to the CTCF motif). The differential effect of these covariates on the TF enrichment was estimated using the adjusted slope (interaction term between the covariate and TF expression) by computing the model margins and the pairwise comparisons of predictive margins using STATA v17.

Bivariate linear regressions between individual TF motif enrichment and expression level were performed using R.

#### AlphaFold3 prediction

For each transcription factor (TF) — KLF4, OCT4, SOX2, MYC, and YY1 — we randomly selected loci for each category of “CTCF-only,” “CTCF-TF cobound,” and “TF-only” sites. The DNA sequences (positive and negative strands) of these loci, centered on the CTCF or TF motif and including 35 bp flanking sequences, were extracted using Bedtools ^41^.

TF and CTCF ZF1-11 amino acid sequences were retrieved from UniProt ^54^: Q61164 for CTCF, Q00899 for YY1, Q60793 for KLF4, P20263 for OCT4, P48432 for SOX2, P01108 for MYC, and P28574 for MAX. The CTCF ZF1-11 positions were obtained from Yang *et al.* ^23^.

AlphaFold3 predictions ^55^ were carried out using the DNA sequences, the CTCF sequence, with or without the TF sequence. The predicted protein-DNA structures were visualized in PyMOL 3.10, and the root mean square deviation (RMSD) between the structures was calculated using PyMOL 3.10.

#### Allele-specific binding analyses

Allele-specific binding (ASB) was performed by separating the ChIP-seq or ATAC-seq reads overlapping phased heterozygous SNPs in the GM12878 cell line using R. Allele-specific BAM and bigwig files were then generated using samtools ^48^ and bamCoverage ^45^. To identify ASB

SNPs disrupting the TF motif, allele-specific motif affinity was computed using the R package, atSNP ^56^.

#### Nano-NOMe-seq single molecules clustering, trajectory and phasing score

For each condition (untreated, IAA) and genomic context (based on accessibility, CTCF signal strength, presence of TF motifs), 5000 molecules were randomly sampled. The level of 5mC methylation in GpC context was binned by 50 bp for each molecule. The missing values were imputed using k-Nearest Neighbors before performing k-means clustering. UMAP was applied using the uwot R package with n_neighbors = 15 and min_dist = 0.2 to project the nucleosome occupancy profiles into two dimensions. For SM trajectories, a principal curve was first fitted on the UMAP-embedded molecules using the princurve R package ^57^. All molecules were then projected onto this curve to obtain pseudotime estimate and trajectory inferred using the slingshot R package ^24^, specifying cluster 1 as the root. Nucleosome phasing scores were calculated by applying a fast Fourier transform (FFT) to the binarized methylation patterns of each read. Two separate phasing scores were computed for accessible and inaccessible sites, targeting periodicities of 190 bp and 170 bp, respectively. For each, the power spectrum was computed and the phasing score was defined as the proportion of signal intensity within a narrow frequency band around the expected nucleosome spacing.

## Data availability

All raw and processed sequencing data files are deposited at NCBI’s Gene Expression Omnibus (GEO) and will be available to public on publication of the manuscript.

## Code availability

No custom code/software was used in the study. The publicly available software used is indicated in the methods section.

## Acknowledgements

This work was supported by 1R35GM122515 (J.S) and NIH P01CA229086 (J.S). GC were supported by a fellowship from the NCC.

The authors thank Skok lab members for helpful scientific discussions, New York University School of Medicine High Performance Computing Facility (HPCF) for computing technical support, Adriana Heguy and the Genome Technology Center (GTC) core for sequencing efforts, NYU Flow Cytometry and Cell Sorting Center for FACS analysis and sorting. GTC is a shared resource partially supported by the Cancer Center Support Grant P30CA016087 at the Laura and Isaac Perlmutter Cancer Center.

The authors thank Sofia Battaglia from the Bernstein’s lab for her advice and protocols for the Nano-NOMe-seq library preparation.

## Author contributions

These studies were designed by Jane Skok and Catherine Do. The Nano-NOME-seq experiments were performed by Guimei Jiang and Paul Zappile under the supervision of Adriana Heguy. The analysis was performed by Catherine Do The paper was written by Jane Skok and Catherine Do.

## SUPPLEMENTARY INFORMATION

### Supplemental Tables

**Table S1. Tab A**: Accession numbers of the CTCF ChIP-seq, ATAC-seq and RNA-seq for human ENCODE samples used in the motif analyses. Only samples with data in all three assays and with at least 35% of CTCF motif-containing peaks were included. The percentage of CTCF motif found using FIMO are listed. **Tab B**: Number and percentage of accessible CTCF peaks in human and mouse including the CTCF ChIP-seq and ATAC-seq accession numbers for the mouse ENCODE samples. This data was used in **Figure S1B.** This table relates to **Figure 1**.

**Table S2. Tab A:** Proportions and counts of the sequences with enriched TF motifs within 35 bp of the CTCF consensus motif at accessible and inaccessible CBSs in WT mESCs. **Tab B:** Proportions and counts of the sequences with enriched TF motifs within 35 bp of the CTCF consensus motif at accessible and inaccessible binding sites in human ENCODE samples with at least 20% of CTCF motif-containing peaks. This table relates to **Figure 1**.

**Table S3.** ENCODE accession numbers of the TF ChIP-seq narrow peak and BAM files in GM12878 to assess the overall percentage of cobound CTCF and TF sites. This table relates to **Figure 3**.

**Table S4.** Allele-specific binding of CTCF associated with allele-specific of TF with SNPs disrupting either CTCF or the TF motifs. This tables relates to **Figure 3**.

**Table S5**. Significant correlations between the percentage of TF motifs within 35 bp of CTCF consensus motif and their expression level in ENCODE human samples. This relates to **Figure 4**.

## Supplemental Figures

**Figure S1.**
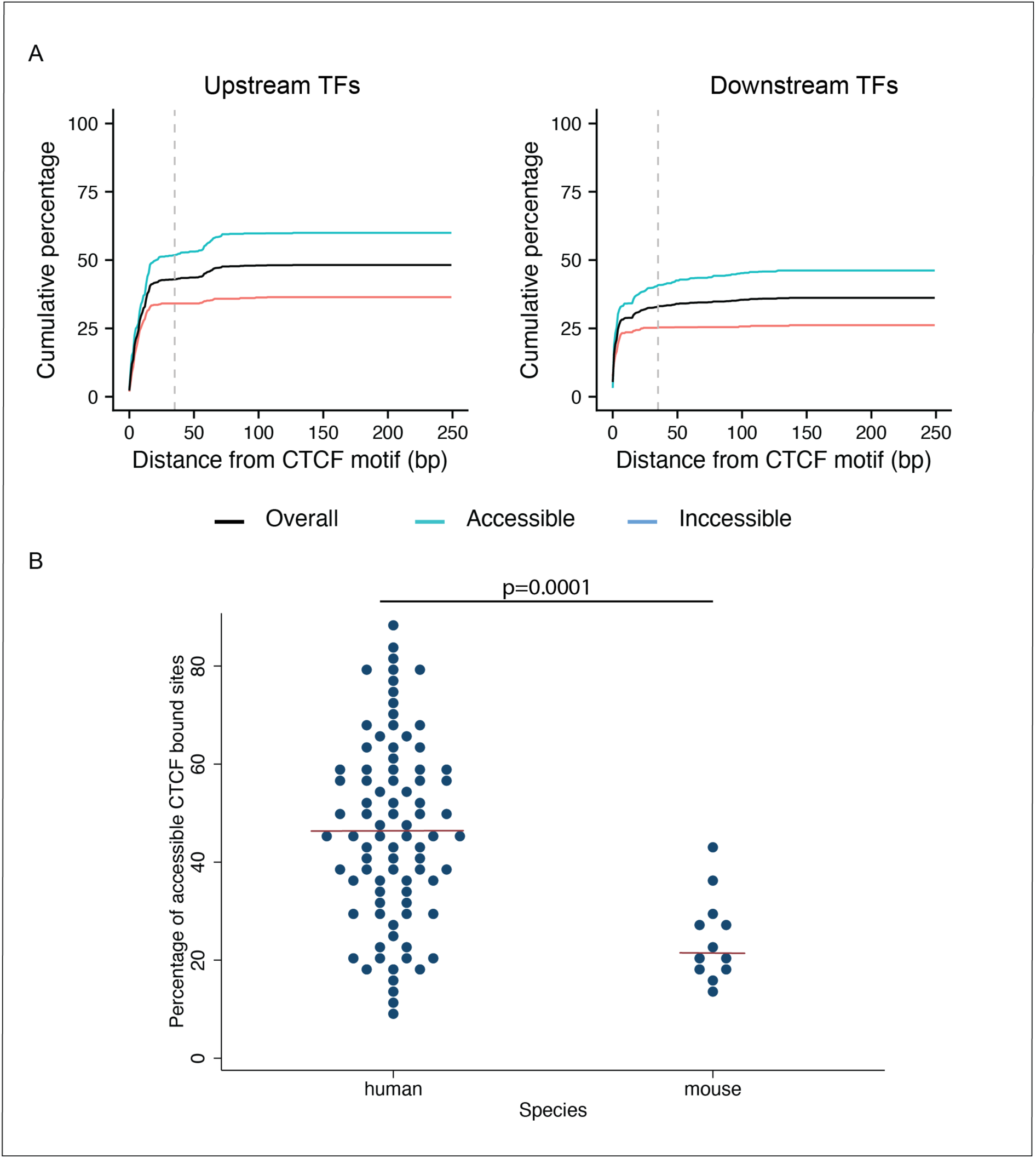
A: Cumulative percentage of CTCF peaks containing a TF motif within 250 bp at accessible and inaccessible sites. The dashed lines correspond to 35 bp upstream or downstream of the CTCF motif. This figure relates to Figure 1. **B:** Dotplot showing the higher percentage of accessible CTCF bound sites in human versus mouse. The red line represents the median. The p-value has been computed using a t-test. This figure is related to Figure 1.

**Figure S2.**
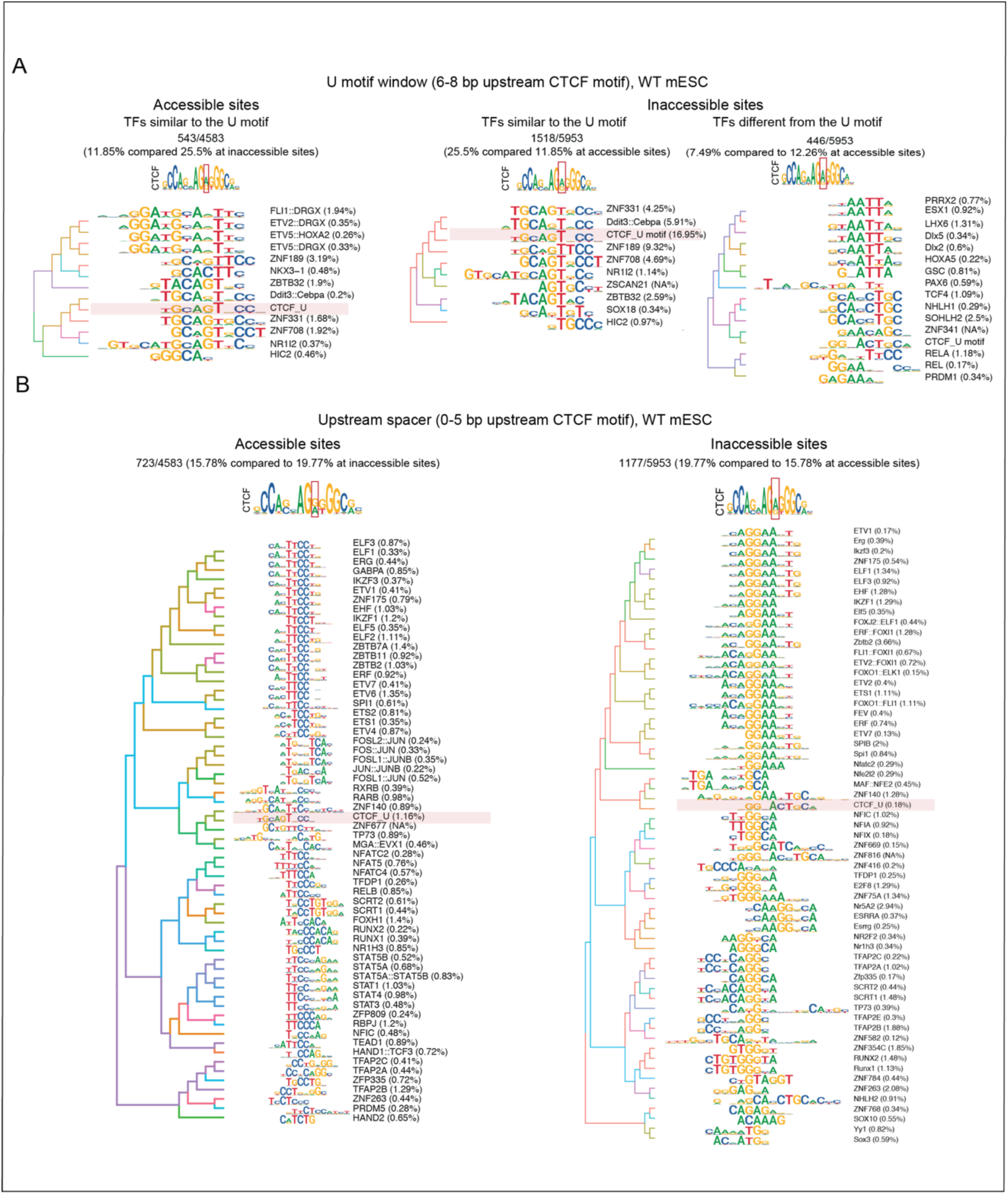
**A**: Motif logos of the enriched TF motifs found in the U motif window at accessible and inaccessible sites in WT mESCs. **B**: Motif logos of the enriched TF motifs found in the U spacer window at accessible and inaccessible sites in WT mESCs. This window is enriched in ETS-family motifs. This figure relates to Figure 1.

**Figure S3.**
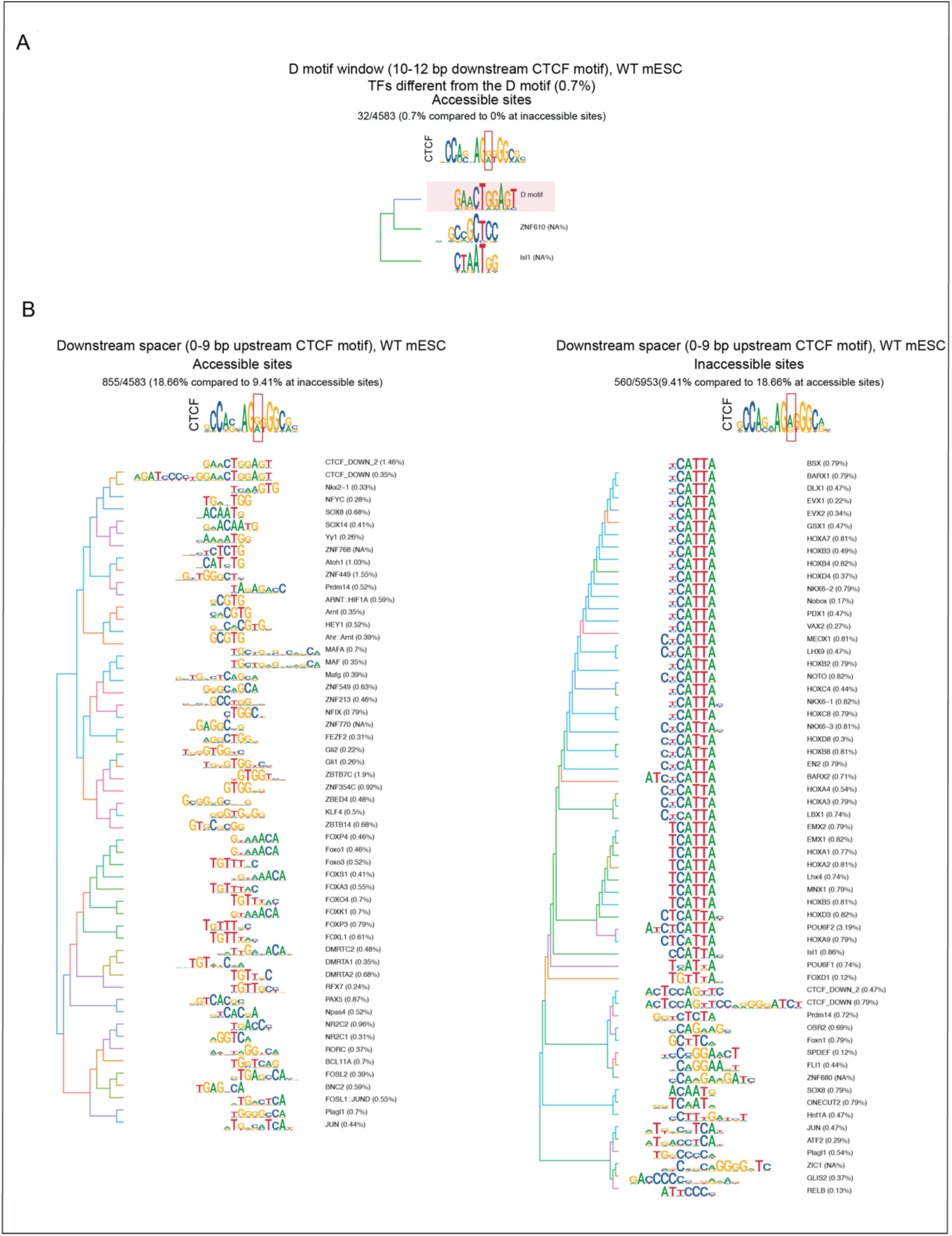
**A**: Motif logos of the enriched TF motifs found in the D window at accessible sites in WT mESCs. No motifs were significantly enriched at inaccessible sites. **B**: Motif logos of the enriched TF motifs found in the downstream spacer window at accessible and inaccessible sites in WT mESCs. This figure relates to Figure 1.

**Figure S4.**
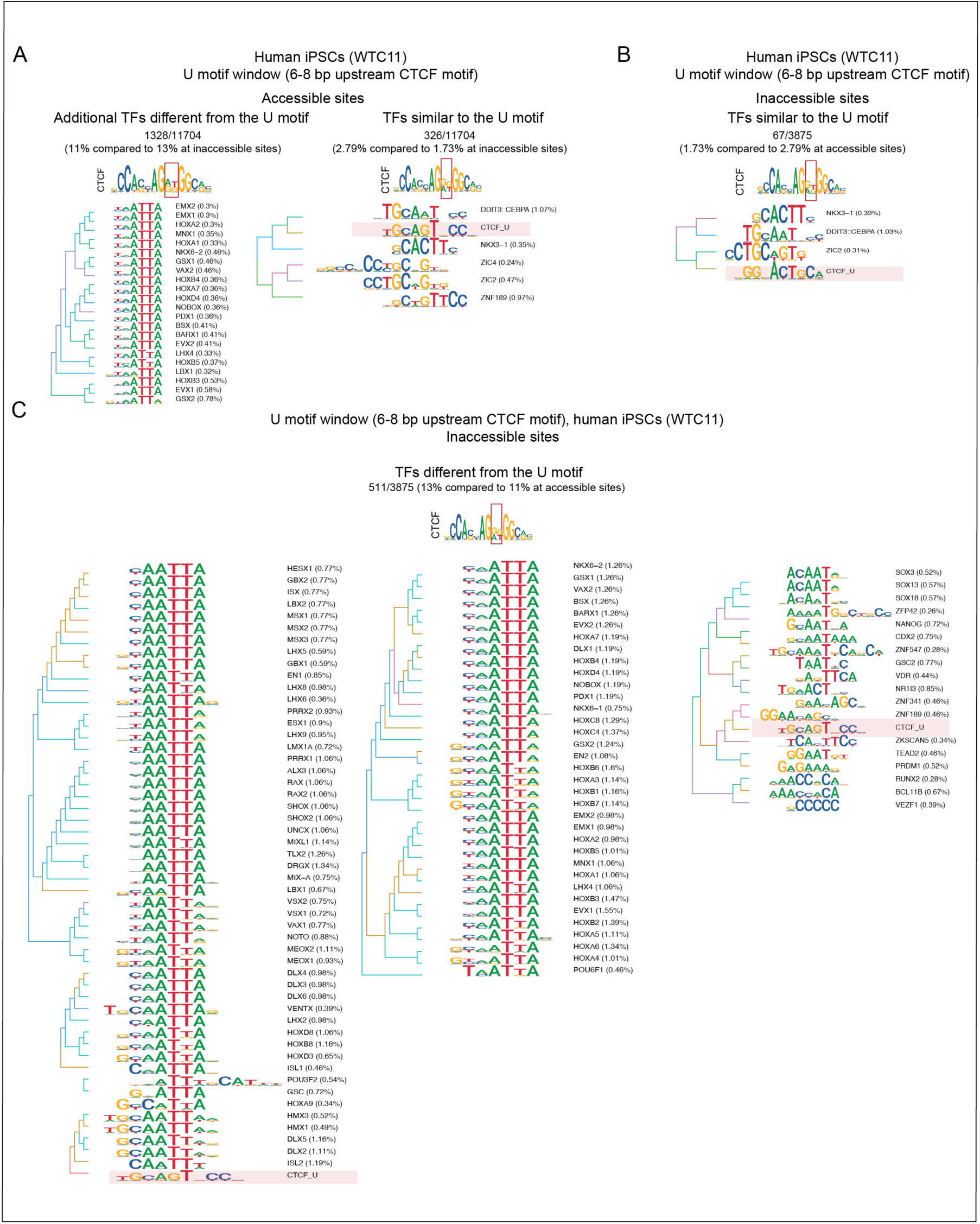
A: Motif logos of the enriched TF motifs found in the U motif window at accessible sites in human iPSCs (WCT11)**. B:** Motif logos of the enriched TF motifs found in the U motif window and resembling the U motif at inaccessible sites in human iPSCs (WCT11)**. C**: Motif logos of the enriched TF motifs found in the U motif window and distinct from the U motif at inaccessible sites in human iPSCs (WCT11). This figure relates to Figure 1.

**Figure S5.**
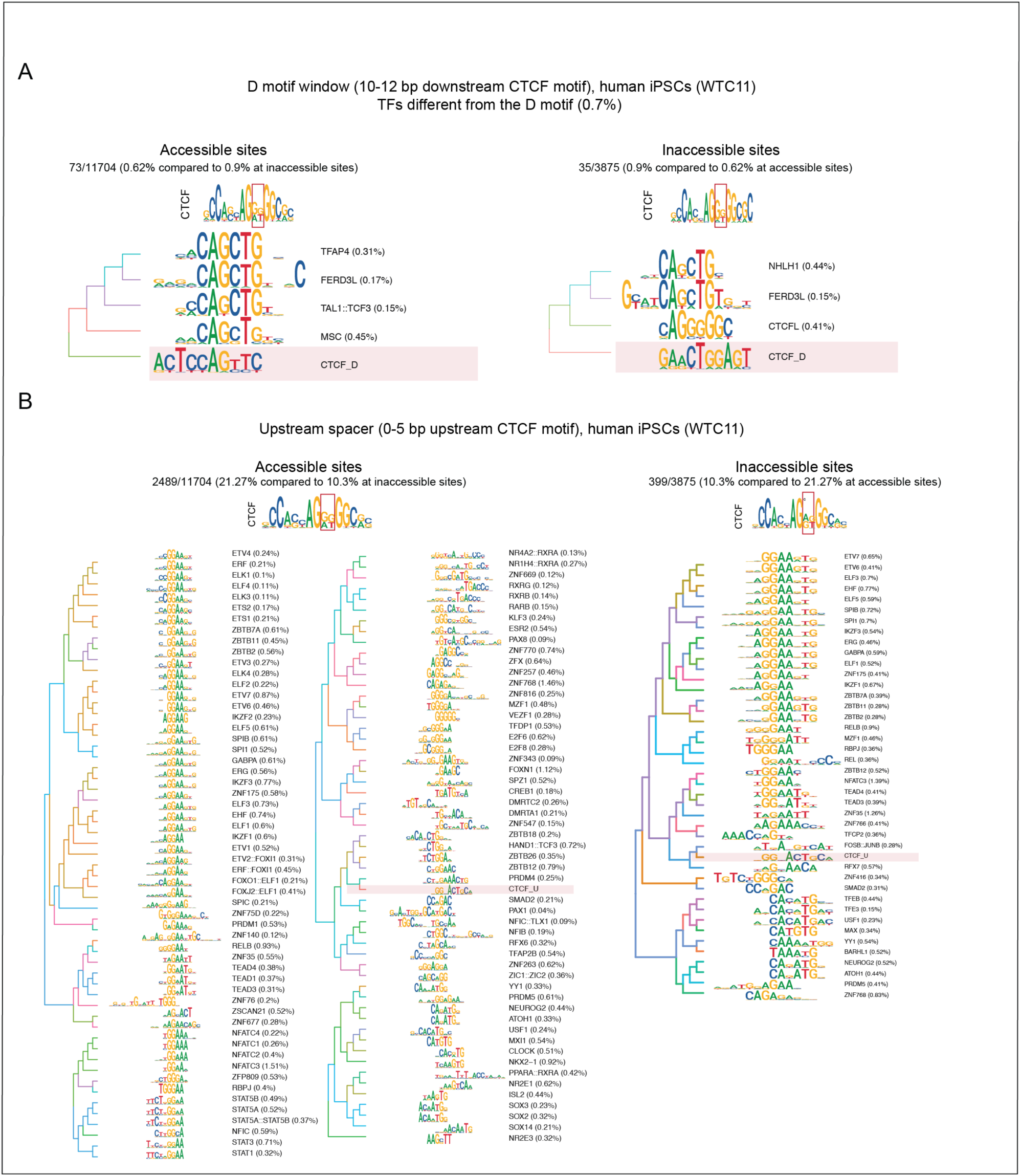
A: Motif logos of the enriched TF motifs found in the D motif window at accessible and inaccessible sites in human iPSCs (WCT11). **B**: Motif logos of the enriched TF motifs found in the upstream spacer at accessible and inaccessible sites in human iPSCs (WCT11). This window is enriched in ETS-family motifs. This figure relates to Figure 1.

**Figure S6.**
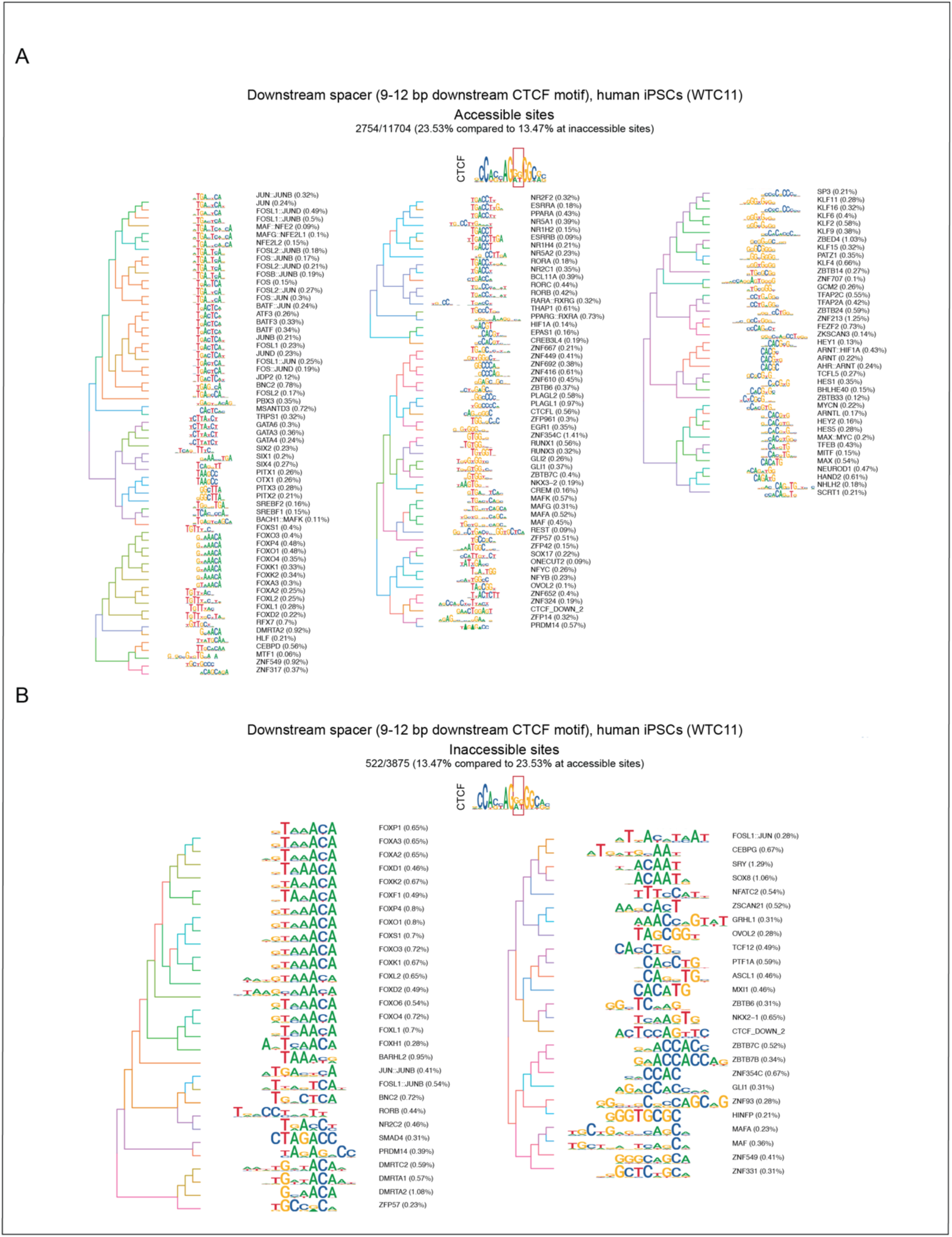
**A**: Motif logos of the enriched TF motifs found in the downstream spacer at accessible and inaccessible (**B**) sites in human iPSCs (WCT11). This figure relates to Figure 1.

**Figure S7:**
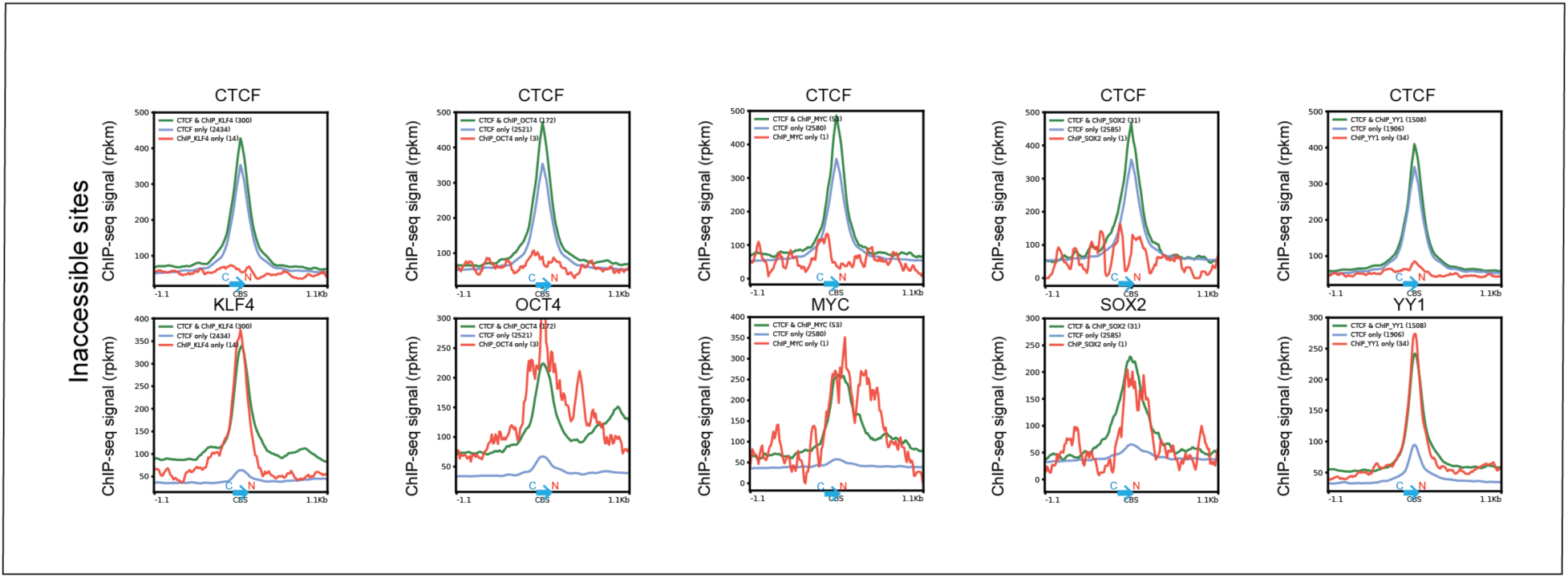
Individual TFs strengthen CTCF signal. Profiles of CTCF (FLAG) and 5 TFs (including 4 pioneer TFs, KLF4, OCT4, MYC and SOX2) showing the effect of TF binding on CTCF signal strength (top) and TF signal strength (bottom) at inaccessible sites where they are co-bound (green) compared to CTCF only (blue) and TF only sites (red). This analysis is restricted to CTCF binding sites with the consensus CTCF motif. The graphs are oriented on the C to N terminal CTCF core motif. This figure relates to Figure 2.

**Figure S8:**
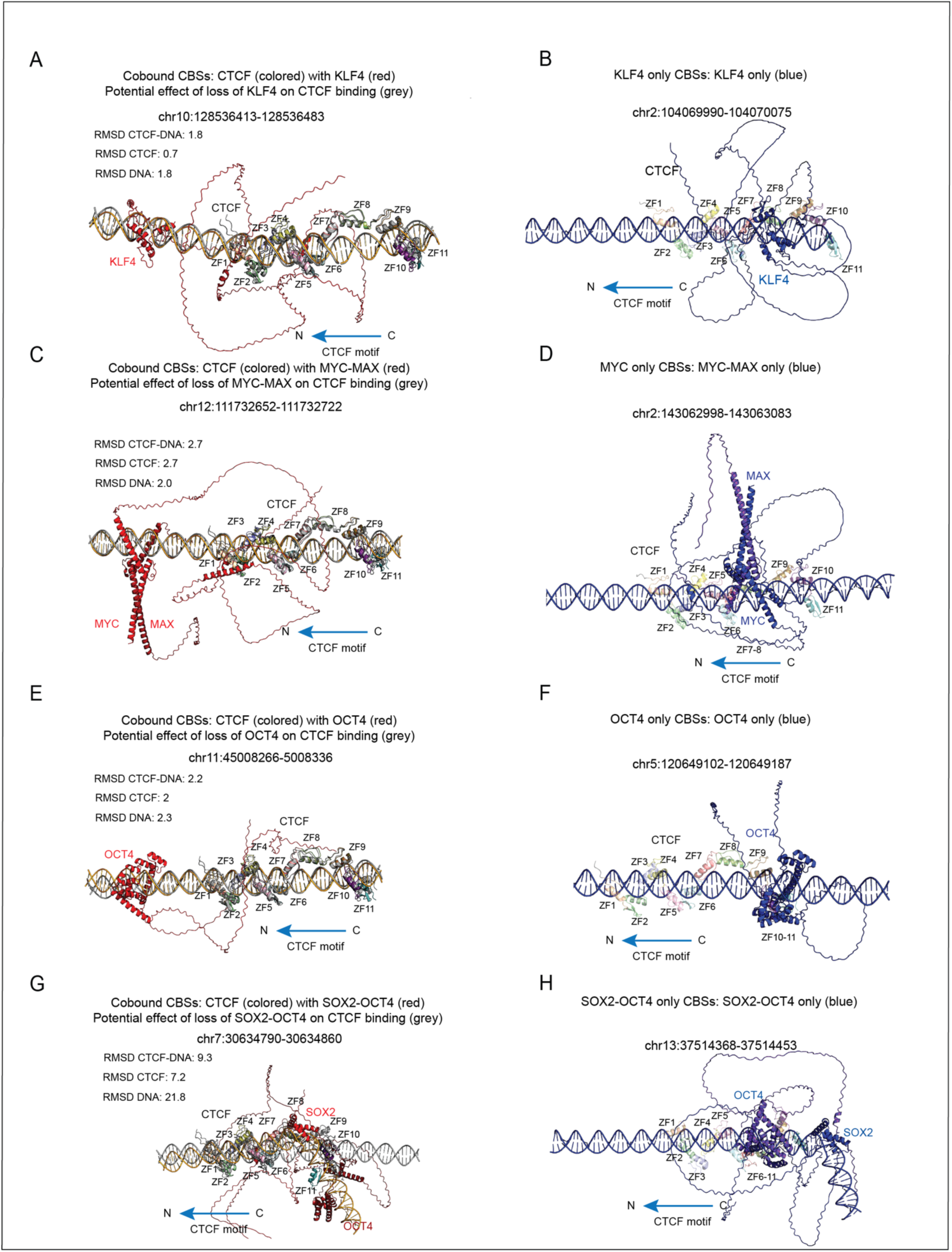
TFs can cobind or compete with CTCF at CBSs. AlphaFold3 predictions at cobound sites for KLF4 (**A**), MYC-MAX (**C**), OCT4 (**E**) and SOX2-OCT4 (**G**). To assess the effect of the cobound TF, the structures of CTCF with (colored) or without TF (grey) were aligned and the RMSD (root mean standard deviation) calculated for the CTCF-DNA complex and separately for CTCF and DNA (top). At cobound sites, TFs tend to bind outside CBSs or in the U spacer (between ZF7 and ZF9). SOX2 binding can induce a major bend in the DNA (**G**). AlphaFold3 predictions at TF only bound CBSs for KLF4 (**B**), MYC-MAX (**D**), OCT4 (without SOX2) (**F**) and SOX2-OCT4 (most of the SOX2 peaks overlap with OCT4) (**H**). TFs are shown in in blue. CTCF binding predictions are overlayed to visualize the position of the TF relative to CTCF. At those sites, the TFs tend to bind at CTCF core or extended binding sites and as a result they directly compete with its binding. This figure relates to Figure 2.

**Figure S9:**
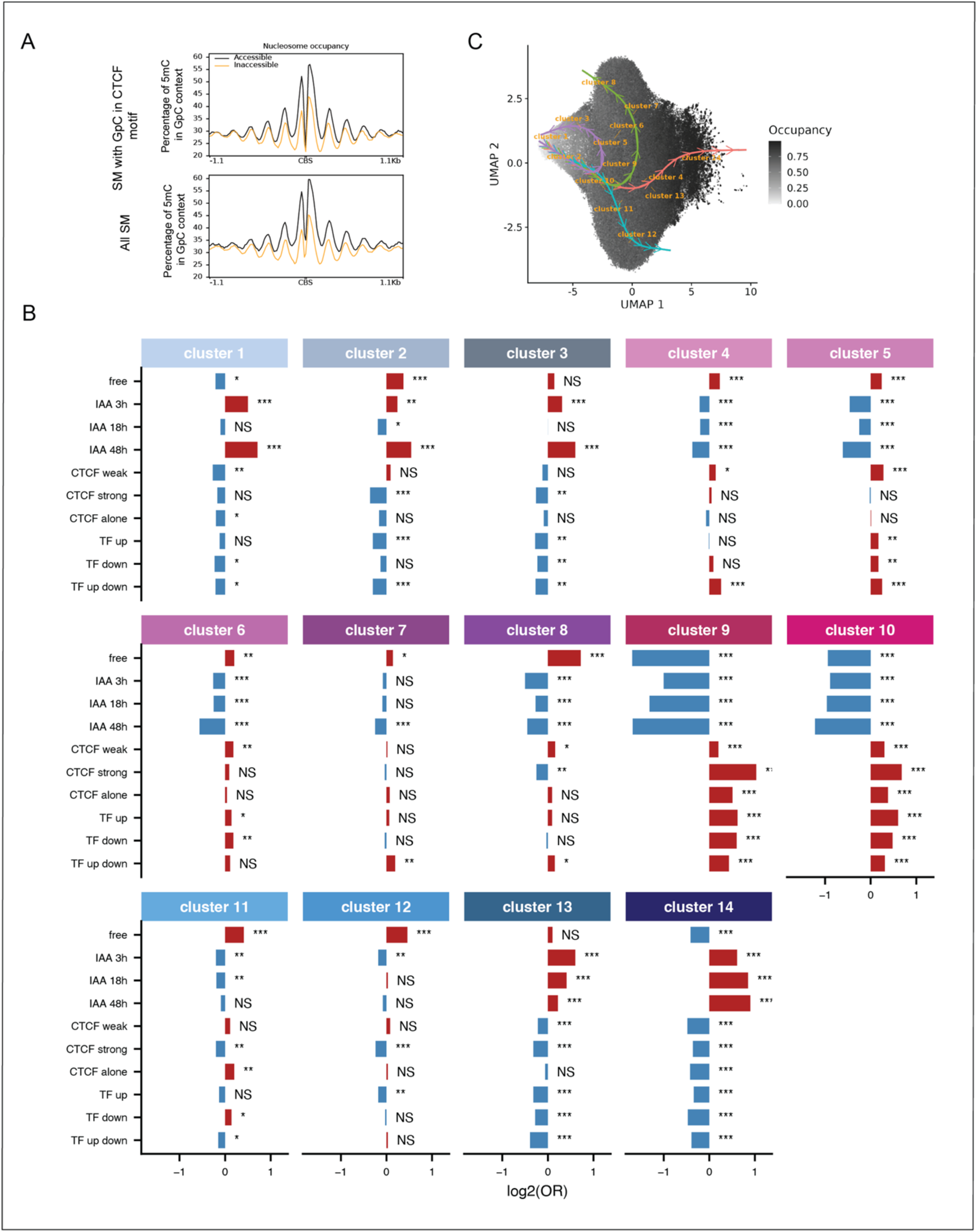
**A**: Profiles showing similar nucleosome occupancy phasing of occupied SM for GpC-containing CBSs compared to non GpC-containing CBSs. **B**: Enrichment of each SM cluster for the condition and genomic context under study. The odd ratios and p-values were computed using logistic regressions**. C:** UMAP of SMs colored by the molecule overall nucleosome occupancy. This figure relates to Figure 5.

**Figure S10:**
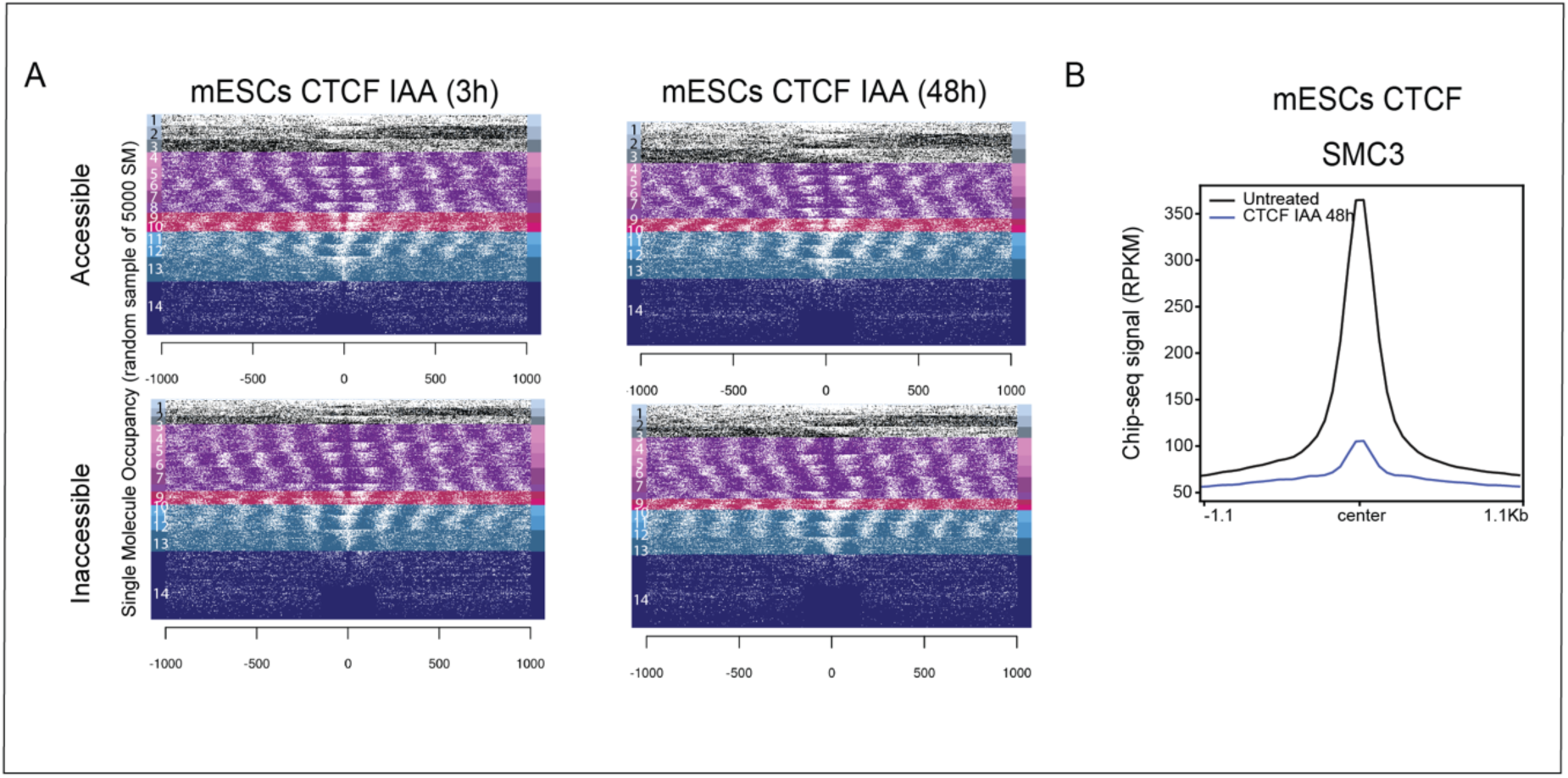
**A**: Heatmaps showing the effect of CTCF degradation on nucleosome phasing after 3h and 48h of IAA treatment in mESCs. **B:** Profiles of SMC3 at lost CTCF peaks after CTCF degradation (48h IAA) at accessible and inaccessible sites in mESCs. This figure relates to Figure 6.

